# Mechanism of bacterial adhesion and embedment in a DNA biofilm matrix: Evidence that binding of outer membrane lipopolysaccharide (LPS) to HU is key

**DOI:** 10.1101/2020.10.20.346890

**Authors:** Bhishem Thakur, Kanika Arora, Archit Gupta, Purnananda Guptasarma

## Abstract

In biofilms, bacteria are embedded within a matrix of extracellular DNA (e-DNA). Since bacterial cells and e-DNA are both negatively-charged, a positively-charged substance must act like a ‘glue’ to allow bacteria to be embedded within the DNA matrix. Here we show that HU (a highly-abundant, histone-like, nucleoid-associated, DNA-binding protein) facilitates bacterium-bacterium and bacterium-DNA interactions by binding to lipopolysaccharide (LPS), a bacterial outer membrane component. We demonstrate that LPS binds to both the canonical and non-canonical DNA-binding sites on HU. We propose that the hexose sugar-terminal phosphate moieties present in the lipid A head-group of LPS bind to the same lysine/arginine residues that are involved in binding of the pentose sugar-phosphate groups in DNA. Alternate binding of LPS and DNA by HU’s DNA-binding sites could allow HU to bind to bacterial cells surfaces and thus elicit bacterium-bacterium and bacterium-DNA interactions in biofilms.

## Introduction

Bacteria in biofilms are embedded within a matrix of extracellular polymeric substance (EPS) consisting predominantly of extracellular DNA (e-DNA) derived from the lysis of bacterial cells,^1^ or from cellular secretions.^2^ One practical problem with understanding such embedment is that negative charges decorate the surfaces of both bacteria and DNA, creating scope for repulsive interactions.^3^ In principle, any protein decorated with positive charges, such as an abundant and non-sequence-specific DNA-binding protein (e.g., a protein involved in the compaction of DNA within bacterial chromosomes) could potentially bind to both DNA and to bacteria, to function as a charge-neutralizing glue.^4^ A good candidate for such a protein glue in bacteria is the highly-abundant, nucleoid-associated, DNA-binding protein, HU, since it is already known to be present in bacterial biofilms (in association with e-DNA derived from bacterial chromosomal DNA).^5^ Importantly, HU is also now known to be limiting for biofilm formation, since anti-HU antibodies have been demonstrated to disrupt biofilms.^6^ In this article, we show that HU binds to free lipopolysaccharide (f-LPS) as well as to the surfaces of bacterial cells [which contain cell-anchored LPS (c-LPS) in their outer membranes]. We also show that that the binding of HU to either f-LPS, or c-LPS, can involve either (i) HU’s canonical DNA-binding site (i.e., one or both of a pair of extended lysine-rich hairpin loops in the HU dimer, one derived from each monomer), or (ii) HU’s non-canonical DNA-binding site (i.e., one or both of a pair of double-lysine clusters on the sides of the HU dimer, one derived from each monomer). We show that addition of micellar f-LPS to free HU (f-HU) generates large molecular assemblies, and that addition of f-HU to c-LPS generates large cellular assemblies (bacterial clumps). The charged head-group present in the lipid A component of LPS contains two hexose sugar-linked terminal phosphate moieties, each bearing two positive charges. We propose that these can bind to lysine/arginine residues that are positioned on (each of) f-HU’s two different types of DNA-binding sites, using specific binding modes, and molecular recognition, i.e., through the presentation of charges to each other under specific distance and geometry restraints.

HU is a bacterial chromosomal nucleoid-associated protein (NAP) of the DNABII class. The reason that it may be a good candidate protein for functioning as a glue which can neutralize negative charges upon both bacterial surfaces and DNA is that it possesses several distinct characteristics which support the likelihood of its being associated with all types of e-DNA, at all conceivable locations, in any biofilm, while remaining capable of binding to other charged substances: (1) The binding of HU to DNA is entirely non-sequence-specific,^7^ allowing it to be ubiquitous on the surface of all e-DNA; (2) The abundance of HU in bacteria is extremely high, reaching up to 50,000 dimeric assemblies per cell during exponential growth,^8^ allowing it to be present in large numbers on all DNA; (3) HU is naturally released into the extracellular milieu, together with bound cellular DNA being transformed into e-DNA, through lysis,^1^ or explosive cell lysis,^9^ and through secretion,^2, 10^ including secretion involving type IV mechanisms;^11, 12^ and (4) HU is multivalent, in that (a) HU can exist in different types of quaternary structure (i.e., as dimers, tetramers or octamers),^13^ and also because (b) each monomeric chain of HU folds to generate (i) a lysine/arginine-rich beta hairpin which may be called the *canonical DNA-binding site*, which can wrap around DNA’s minor groove,^14^ and (ii) a double-lysine cluster on the side of the HU dimer which may be called the *non-canonical DNA-binding site*, which allows HU to bind to DNA through an additional recently-discovered mode.^15^ In the remainder of this introductory section, we introduce the structural-biochemistry of the HU dimer, and the locations of its canonical/non-canonical DNA-binding sites (Figures 1A–1F).

**Fig. 1.**
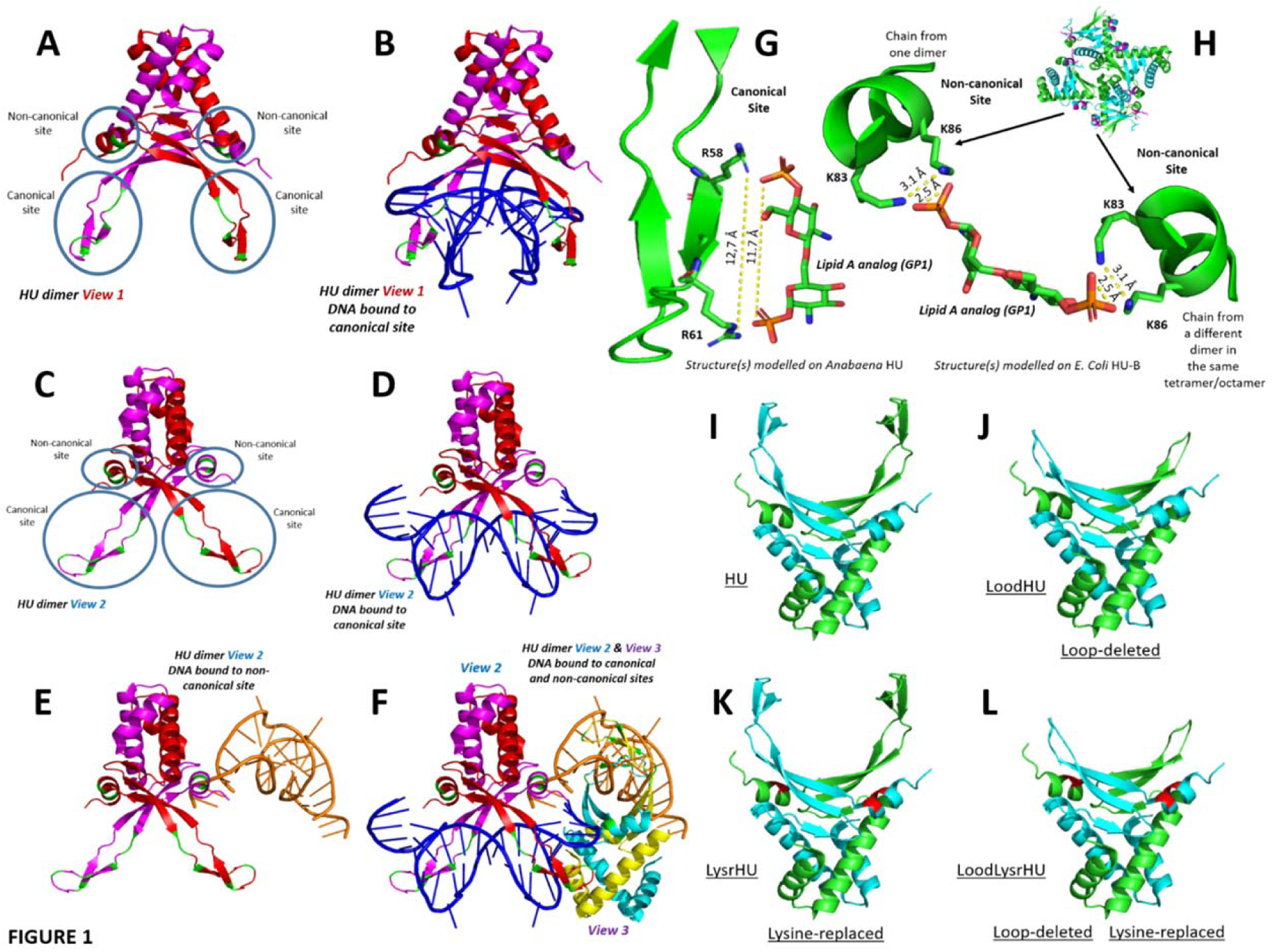
HU’s canonical and non-canonical interactions with DNA and potential modes of interaction with LPS; variants created to probe interactions with DNA and LPS. **a**, An HU dimer (view 1; *Anabaena* HU; derived from PDB ID 1P51) showing locations of the canonical and non-canonical DNA-binding sites. **b**, The same HU shown in panel a (view 1) with a 20-mer double-stranded DNA molecule bound at its canonical DNA-binding site. **c**, The same HU shown in panel a, but rotated by 45 degrees to obtain a different view (view 2). **d**, The same (DNA-bound) HU shown in panel b, but rotated by 45 degrees (view 2), clearly showing the beta hairpin loops from each of the two monomers grasping DNA around the minor groove. **e**. The same HU shown in panel c, but with the canonical site unoccupied and with a 20-mer double-stranded DNA molecule bound at the non-canonical DNA binding site. **f**, The same HU shown in panels c, d and e, but with both canonical and non-canonical DNA binding sites occupied by 20-mer double-stranded DNA molecules; in this arrangement, two HU dimers from a single asymmetric unit of the crystal are shown (one from view 2 and the other from a different view, view 3); each dimeric HU’s canonically-bound DNA is also bound to the non-canonical site of the other dimeric HU. **g**, A proposal based on the structure of *Anabaena* HU (PDB ID 1P51) regarding use of two conserved arginine residues (R58 and R61) on a beta hairpin loop from the canonical DNA-binding site for binding to the sugar-phosphate head group of the lipid A component of lipopolysachharide (LPS); here, a stick representation of the epsilon amino groups of arginine is used to establish distance compatibility with the two terminal phosphate groups in the head group of lipid A. **h**, A proposal based on the structure of *E. coli* HU (PDB ID 2O97) regarding the use of two conserved lysine residues (K83 and K86) on a helix on the side of each monomer of HU for binding to a doubly-charged terminal phosphate in the sugar-phosphate head group of the lipid A component of LPS, with the other terminal phosphate bound to a different helix from a different dimer in an HU higher-order multimer. **i**, Wild type HU with both canonical and non-canonical DNA binding sites present. **j**, A schematic of loop-deleted HU (LoodHU) in which the canonical DNA-binding sites are deleted and replaced with an 11 amino acids-long glycine/serine-rich linker. **k**, A schematic of lysine-replaced HU (LysrHU) in which lysines K83 and K86 at the non-canonical DNA-binding sites (shown in red) are replaced by alanine. **l**, A schematic of loop-deleted and lysine-replaced HU (LoodLysrHU), in which both canonical and non-canonical DNA-binding sites are missing. The PDB ID 1P51 was used to generate the schematics in panels i-l.

*Locations of canonical and non-canonical DNA-binding sites upon HU’s structure.* Figure 1A highlights, in green, the polypeptide backbone locations of the two canonical DNA binding site(s) of HU, i.e., one site present per monomeric HU chain. These sites are shown upon the structure of the *Anabaena* HU dimer, as identified by X-ray crystallography;^14^ the figure also shows where the non-canonical DNA-binding site(s) identified later (using small-angle X-ray scattering studies of the structurally-analogous *E. coli* HU molecule),^15^ physically locate upon this structure. The reason that Figure 1A shows *Anabaena* HU instead of *E. coli* HU (whose X-ray crystallographic structure is also known),^16^ is that no electron density is seen in the latter for the canonical DNA-binding loops in the absence of DNA, presumably owing to mobility of these loops. Figure 1B shows the *Anabaena* HU structure in complex with a double-stranded 20-mer fragment of DNA (in blue) bound at the canonical DNA-binding site. Figures 1A and 1B represent HU from the same viewing angle (View 1). They are derived from a crystal in which two dimers of *Anabaena* HU display crystallographic association; basically, each 20-mer (HU-bound) DNA fragment is also bound to a different HU dimer within the asymmetric unit, through the terminal regions of the DNA fragment.

In the remaining panels of the Figure 1, we use these additional noncovalent interactions to schematically illustrate how HU can simultaneously bind to DNA through both its canonical and non-canonical DNA-binding sites, in principle. Firstly, in Figure 1C, we show the HU dimer from Figure 1A in a different view (View 2) rotated by 45 degrees in respect of View 1, along a vertical axis. Correspondingly, Figure 1D shows the DNA-bound view of the rotated HU dimer in Figure 1C. Figures 1E and 1F display interactions of HU with DNA at the non-canonical site. Using View 2, in Figure 1E, a (second) different copy of the double-stranded 20-mer DNA present in the same asymmetric unit (bound to the canonical site of a different dimer) is shown interacting with the non-canonical site of the dimer shown in Figures 1A–1D (in yellow). Finally, in Figure 1F, a complete view of both dimers in the asymmetric unit is shown, with each dimer associating with one copy of the 20-mer double-stranded DNA at its canonical site, and with a different copy of the 20-mer double-stranded DNA at its non-canonical site. In Figure 1F, the second HU dimer is shown from a third view (View 3) which is different from Views 1 and 2. Figure 1 may thus be taken to be a schematic diagram; one that uses real crystal structures and interactions of *Anabaena* HU with DNA to illustrate how the analogous *E. coli* HU might potentially simultaneously interact with DNA at both its canonical site and also at the non-canonical site recently identified through SAXS studies.^15^

## Results and Discussion

### Section 1: Structural analyses and considerations

#### Analyses of possible binding geometries of LPS to HU’s canonical and non-canonical DNA-binding sites

Based on the structural-biochemical perspectives introduced towards the end of the above section, we decided to first theoretically examine whether HU’s DNA binding sites could also bind to negatively-charged lipopolysaccharide (LPS) molecules present on the outer membranes of bacteria, i.e., either to LPS in the form of free LPS (f-LPS) that has been shed into the extracellular medium, or to cell-displayed LPS (c-LPS). We examined distances between negatively-charged phosphate moieties in LPS and positively-charged lysine/arginine residues present at the DNA-binding sites. The head-group of the lipid A component of LPS contains two negatively-charged hexose-linked phosphate moieties. There are two important differences between these moieties and the analogous sugar-phosphate moieties in DNA. Firstly, the sugars in DNA/RNA are pentose sugars (deoxyribose or ribose) whereas those in LPS are hexose sugars. Secondly, the phosphate groups in the DNA backbone contain only one negative charge per phosphate due to the participation of the phosphate in the formation of phosphodiester bonds; in contrast, in the head-group of lipid A, both phosphates are terminal phosphates and, therefore, possess two negatively charged oxygen atoms each. Notably, the phosphates on the head-group of the lipid A component of LPS are already known to bind to poly-L-lysine through electrostatic interactions.^17^ Based on similar interactions, Figures 1G and 1H shows schematically how these phosphates in lipid A can also potentially bind to lysine/arginine groups present at the DNA-binding sites of HU, using the structures of the polypeptide backbones for segments of the canonical (Figure 1G) and non-canonical (Figure 1H) DNA-binding sites of *Anabaena* HU in which the side chains of the relevant lysine and arginine residues have been shown in green (stick model). The backbones of the HU segmental structures have been shown juxtaposed against the known crystal structure of the head-group of lipid A (which was sourced from the crystal structure of an antibody bound to the same head-group).^18^ Figure 1G shows how the two phosphates groups in the lipid A component of LPS, one each at the two ends of the head-group, can potentially bind to two different ε-amino groups (separated by 12.7 Å) located upon arginine residues, R58 and R61, in HU’s canonical DNA-binding site. These two arginine residues are conserved between *Anabaena* and *E. coli* HU and also in HU from most other bacteria.^19^ Figure 1H shows how the two negative charges present upon the terminal phosphate group present at any end of the head-group of lipid A could potentially bind to two ε-amino groups (separated by 3.1 Å) located upon two lysine residues, K83 and K86, on the same face of a helix in HU’s non-canonical DNA-binding site. Together, Figures 1G and 1H suggest that a theoretical case exists for LPS to bind to HU, not merely through electrostatic interactions but through specific, electrostatic interactions between phosphate groups and lysine/arginine residues with possible distance restraints suggestive of molecular recognition.

#### Analyses of variants of HU that could be created to examine DNA and LPS binding to canonical and non-canonical DNA-binding sites

To examine the interactions of DNA and LPS with HU, we created protein-engineered HU variants. Before describing the analyses performed to decide upon the details of these variants, we would like to point out that in *E. coli* HU exists in two isoforms, HU-A and HU-B, which are both ∼90 amino acids long,^20^ and share approximately ∼69 % sequence identity. HU-A, is synthesized mainly in the late lag and early log phase of growth and exists predominantly as a dimer or a tetramer, whereas HU-B is synthesized mainly in the mid-log phase, and exists as dimers, tetramers or octamers.^13, 16, 21^ In the late-log phase and stationary phase, it is believed that heterodimers of HU-A and HU-B exist in combination with DNA.^22^ In a previous paper, we have described engineered forms of HU-A and HU-B, in which each protein has been genetically fused with a fluorescent protein at its N-terminus. We had created chimeric forms in which a monomeric red fluorescent protein (tag-RFP) was fused with HU-A (to make RFP-HU-A).^23^ Similarly, a monomeric yellow fluorescent protein (Venus) was fused with HU-B (to make Venus-HU-B). In the work described here, we have used both 6xHis-tagged and affinity-purified wild-type forms of HU-A and HU-B, without any fluorescent protein in fusion, and also the engineered forms referred to above (i.e., RFP-HU-A and Venus-HU-B), in microscopic, cytometric and other investigations aimed at discovering whether f-LPS or c-LPS can bind to f-HU, RFP-HU, or Venus-HU. Further, we also constructed variants of HU-A and HU-B called LoodHU, LysrHU and LoodLysrHU, as explained below. As already observed in Figures 1A and 1G, the canonical DNA-binding (and putatively also LPS-binding) site on an HU dimer consists of two loops, one derived from each HU monomer. Likewise, in each monomer as already observed in Figures 1A and 1H, the non-canonical DNA-binding (and putatively also LPS-binding) site consists of a double-lysine cluster in each monomer, consisting of lysine residues K83 and K86. Thus, we created four different variants of HU-A and HU-B: (i) wild-type HU (Figure 1I) containing both canonical and non-canonical DNA-binding sites, (ii) loop-deleted HU, or <u>LoodHU</u> (Figure 1J), in which the 22 residues-long loop (extending from residue 52 to residue 74) is deleted and replaced by an 11 residues-long, glycine/serine-rich linker peptide (N-SGGGGSGGGGS-C), to ablate the canonical DNA-binding loop/site, (iii) lysine-replaced HU, or <u>LysrHU</u> (Figure 1K), in which lysine residues, K83 and K86, are both replaced by the residue, alanine, to ablate the non-canonical DNA-binding site, and (iv) loop-deleted and lysine-replaced HU, or <u>LoodLysrHU</u> (Figure 1L) in which both canonical and non-canonical DNA-binding sites have been ablated through genetic manipulation. Further, we also made variants in which mutations K83A and K86A were made individually.

### Section 2: Binding of HU to free LPS (f-LPS)

#### Microscale Thermophoresis (MST): Binding of f-LPS affects RFP-HU-A diffusion in temperature jump experiments in capillaries

Figures 2A and 2B shows a titration of RFP-HU-A against different concentrations of micellar f-LPS, performed to examine whether binding of f-LPS to HU-A occurs to reduce the rate of diffusion of RFP-HU-A. Here, the diffusion of RFP-HU-A is either away from, or back towards, a region of ‘temperature jump-induced’ molecular depletion within a set of capillaries containing fixed HU and differing LPS concentrations. A dose-dependence of the rate of reduction (through depletion), as well as of the subsequent rate of growth (through replenishment) of the RFP fluorescence signal is observed. This shows that the diffusion of RFP-HU-A, both away from and back towards a region of temperature jump-induced depletion of molecules tends to be progressively slower with greater concentrations of micellar f-LPS. This shows that RFP-HU binds to micellar f-LPS.

**Fig. 2.**
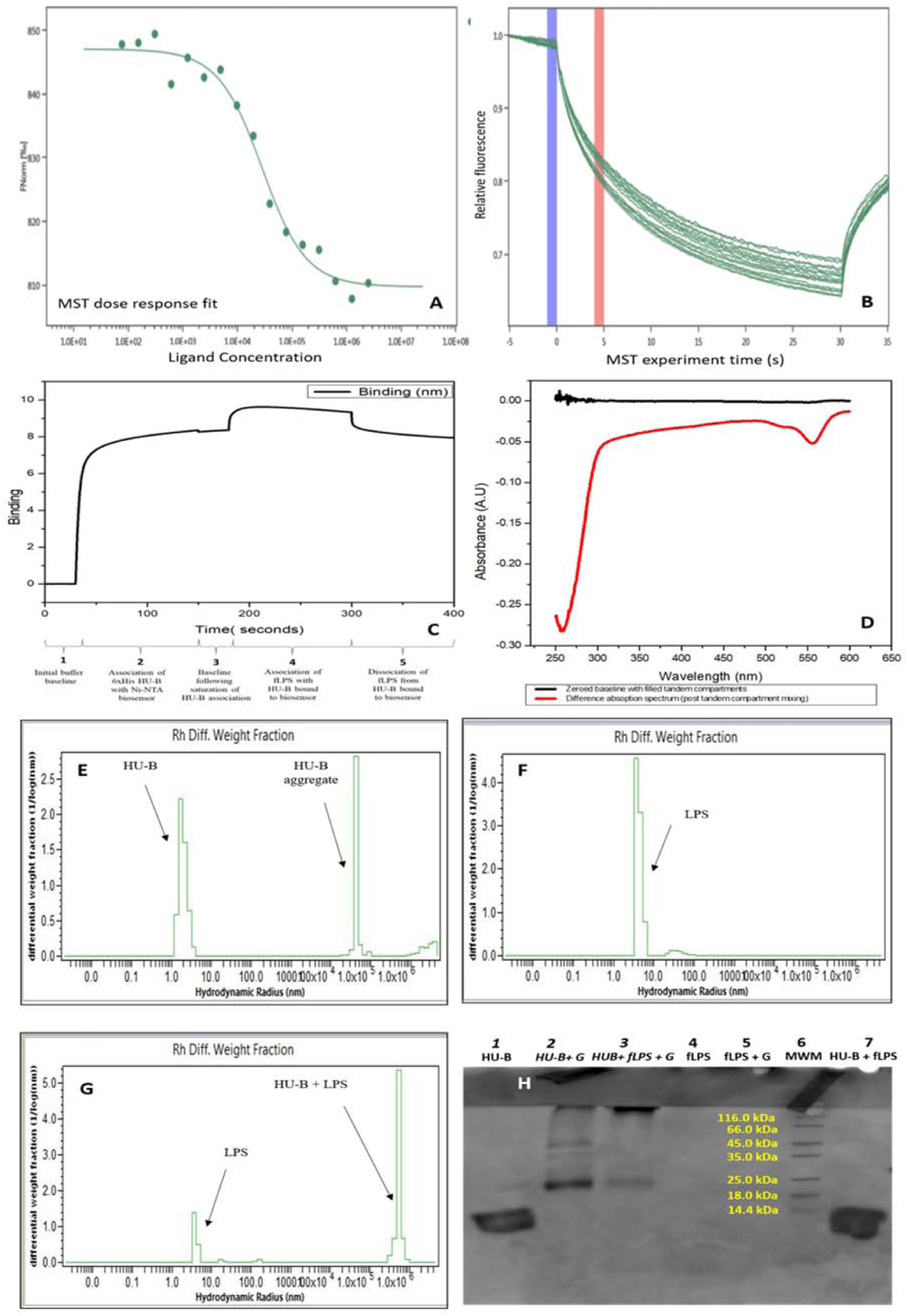
Binding of HU to f-LPS. **a**, Microscale thermophoresis (MST) dose response curve for the binding of f-LPS by HU-A, based on varying concentrations of f-LPS and a fixed concentration of fluorescent tag-RFP-HU-A and plotting rates of reduction in fluorescence from tag-RFP-HU-A due to (temperature jump-dependent) diffusion of molecules away from a microscopic volume of solution being monitored inside a capillary, with different capillaries used for different f-LPS concentrations. **b**, Raw microscale thermophoresis data traces for sixteen capillaries containing different concentrations of f-LPS; blue and red bars indicate the beginning and ending of the time period used to calculate normalized rates of reduction in tag-RFP-HU-A fluorescence for capillaries over a 30 seconds-long laser-induced temperature jump; each curve plots the fall in fluorescence in a separate capillary, and the subsequent rise, over 5 seconds, after heating is switched off. **c**, Bio-layer interferometery (BLI) sensorgram showing the washing baseline (segment 1), binding of HU-B to the Ni-NTA-derivatized sensor (segment 2), washing-baseline (segment 3), binding of f-LPS to HU-B (segment 4) and washing-based dissociation (segment 5). **d**, Difference absorption spectroscopy (DAS) data monitoring binding of HU-B to f-LPS after zeroing of baseline; pre-mixing spectra (black) and post-mixing spectra (red) are used to detect hyper/hypochromic effects in HU-B phenylalanine (∼265 nm) and RFP chromophore (550 nm) absorption bands due to binding. **e, f, and g**, Dynamic light scattering (DLS) monitoring changes in size of 0.02 μm-filtered HU-B, f-LPS and HU-B + f-LPS to detect non-covalent crosslinking of f-LPS by HU-B. **h**, denaturing SDS-PAGE investigating covalent (glutaraldehyde-mediated) crosslinking of HU-B by f-LPS.

The estimated binding constant from these studies can be determined to be ∼34 μΜ, if one assumes a ligand-binding of one LPS molecule per HU chain monomer, and an average molecular weight of 15,000 Daltons for the f-LPS. Of course, both these assumptions are likely to be wrong, given that all f-LPS molecules do not have the exact same molecular weight, and also the fact that the LPS concentrations used here are in the post critical micellar concentration range, as well as the fact that the HU population consists of multiple quaternary structural forms. Therefore, the value of ∼34 μΜ derived in this experiment must only be taken to be indicative of the significantly loose (and not tight) nature of the binding of HU to LPS, and not an accurate measure.

#### Biolayer interferometry (BLI): f-LPS binds to HU-B to generate sensorgrams

Figure 2C shows sensorgrams (similar to sensor-grams in surface plasmon resonance experiments, but detected using a different technique called biolayer interferometry, or BLI) for the binding of 6xHis affinity-tagged HU-B onto a Ni-NTA derivatized probe, and for the further binding of f-LPS in solution to this Ni-NTA-bound 6xHis-tagged HU-B. An association of the HU-B with the probe tip is observed, as is an association of micellar f-LPS with the tip-bound HU-B. The micellar f-LPS is is observed to cause the LPS-HU association curve to descend somewhat even during the association phase, presumable owing to distortions caused by progressively greater binding of micellar f-LPS which could cause the bound molecular masses to shift away from the probe tip’s surface, or even cause some leaching. However, the effect seen during association is small, and a clear dissociation response is also seen to result from the dissociation of f-LPS from HU following depletion of f-LPS from solution. This establishes that HU-B binds to f-LPS.

#### Difference Absorption Spectroscopy (DAS): Binding of f-LPS alters the UV-Visible absorption of RFP-HU-A

Figure 2D shows ‘instrument-zeroed’ absorption profiles of pre-mixing (black) and post-mixing (red) states of an experiment in which equal volumes of RFP-HU-A and f-LPS were mixed. In the present experiment, RFP-HU-A and micellar f-LPS were initially present in the separate compartments of a split-quartz (tandem-compartment) cuvette with a separating wall rising to two-thirds of the cuvette’s height, allowing mixing of the contents of the two compartments to be effected through inversion of the cuvette (after closing of the lid). Thus, mixing of equal volumes of potentially interacting species, followed by refilling of both compartments through ‘up-righting’ of the cuvette, results in halving of the concentrations of the species in each compartment, as well as doubling of the path-length of light passing through solutions of each species, i.e., RFP-HU-A and f-LPS, because after mixing both species fall back into both compartments. A difference in absorbance of light passing through both compartments is anticipated if, and only if, there are interactions between the species in the two compartments, manifesting as a deviation(s) from the zero-baseline, owing to inter-molecular interactions that alter the electronic states of chromophores adjacent to interacting surfaces; otherwise, due to the halving of concentrations and doubling of path-lengths, no difference in absorbance is expected on account of the Beer-Lambert law, as it applies to a DAS experiment.^24, 25^ In the present experiment, mixing is clearly shown to produce deviations manifesting as bands corresponding to reduction in aromatic absorption of HU around ∼260 nm (owing to changes in absorptivity of HU’s phenylalanine residues) and reduction in absorption of the tag-RFP chromophore, at ∼550 nm. These two red bands in the DAS spectrum at ∼260 and ∼550 nm show that RFP-HU-A binds to micellar f-LPS.

#### Dynamic Light Scattering (DLS): HU-B crosslinks non-micellar (filtered) f-LPS into large noncovalent assemblies

Figure 3 shows light scattering profiles which establish that HU-B (with an average size of ∼3-4 nm; Figure 2E) upon addition to f-LPS (with an average size of ∼8–9 nm; Figure 2F), yields large molecular assemblies with sizes in the range of 6×10^6^ nm (an order of magnitude larger than the aggregates of HU, with an average size of 5×10^5^ nm; Figure 2G), with concomitant reduction of the populations corresponding to both HU and f-LPS. Such molecular assemblies are anticipated to form upon the binding of negatively-charged f-LPS to HU-B, since the protein exists as a variety of multimers (dimers, tetramers and octamers) in which each dimer could have two (or more) sites of interaction with negatively-charged LPS. Effectively, HU appears to non-covalently ‘crosslink’ f-LPS into extremely large assemblies. It must be noted that, in these specific (DLS) experiments, unlike in all the other experiments described in this section, the f-LPS is filtered through a 0.02 nm filtration device prior to light scattering with control f-LPS. Thus, we do not see any evidence of the presence of f-LPS micelles (60–95 nm diameter), and are operating in the pre-CMC range of LPS concentrations in which the filtered f-LPS has a diameter of about 8-9 nm, consisting of monomeric LPS or small LPS aggregates. This very clearly establishes that HU-B binds to f-LPS in both micellar and non-micellar forms.

**Fig. 3.**
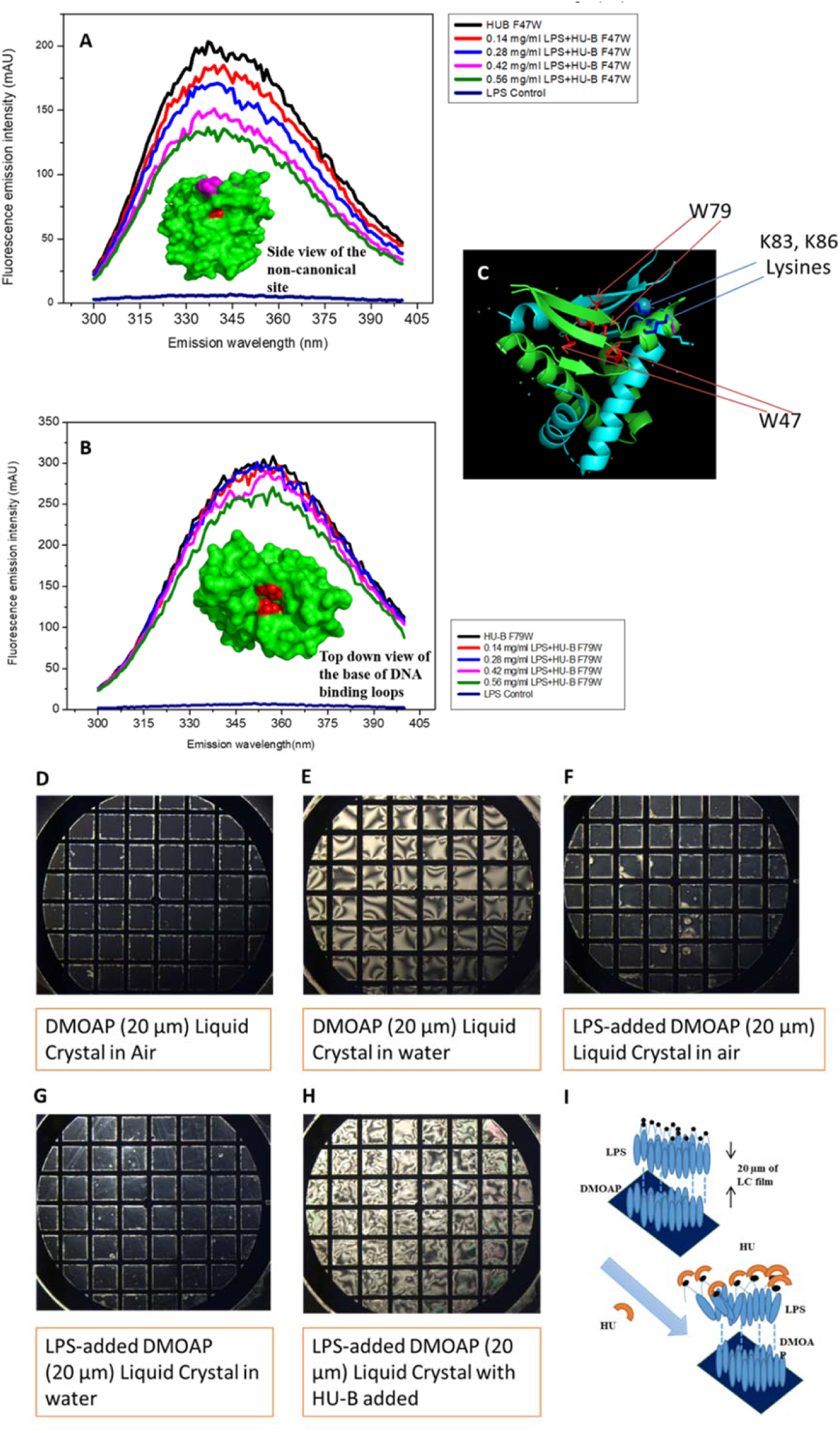
Binding of HU to f-LPS and i-LPS. **a**, Quenching of tryptophan fluorescence in mutant (F47W) HU-B by f-LPS binding; in the inset, residue W47, from one monomer is shown in red, and seen to be located proximal to the non-canonical DNA-binding site residues K83 and K86, shown in magenta. **b**, Quenching of tryptophan fluorescence in mutant (F79W) HU-B by f-LPS binding; in the inset, residue F79, from each of two monomers (facing each other) is shown in red, lying at the base of the canonical DNA-binding site. **c**, A ribbon diagram representation of the structure of the HU-A-HU-B heterodimer (PDB ID 2O97) with superimposed stick representations of residues F47 and F79 (sites of mutations F47W and F79W) as well as residues K83 and K86. Optical polarization micrographs of birefringence from liquid crystals without i-LPS **d**, in air; and **e**, under water; and with i-LPS **f**, in air; and **g**, under water; and **h**, upon addition of HU-B. **i**, A schematic cartoon showing the mechanism of changes in birefringence of the liquid crystal upon interaction of HU-B with i-LPS.

#### Glutaraldehyde addition: Glutaraldehyde cross-links HU-B and micellar f-LPS into large covalent assemblies

Figure 2H shows that upon incubation of HU-B with micellar f-LPS and glutaraldehyde (lane 3), the original HU monomer population in an SDS-PAGE (lane 1) disappears and is replaced by an f-LPS-HU-B molecular assembly that is so large that it is unable to enter into the resolving gel in the SDS-PAGE, after traversing the stacking gel (lane 3). Only a residual faint population of dimeric HU-B is observed. In contrast, glutaraldehyde itself has no comparable effect (lane 2) upon HU alone, i.e., only dimeric and tetrameric populations of HU are stabilized through glutaradehyde crosslinking in the absence of f-LPS, with some crosslinking leading to stabilization of trimers, and with the monomeric band being no longer seen (indicating that all dimers have at least one inter-chain crosslink). Also no band of intensity comparable to that seen in lane 3 is observed in lane 2, at the stacking-resolving gel interface, indicating that glutaradehyde crosslinking of HU-B does not generate large crosslinked assemblies that are comparable to those generated by crosslinking of HU-B to f-LPS. The control in lane 5 shows that no band is seen with addition of glutaraldehyde to f-LPS; obviously, this is because the stain (Coomassie) does not bind to f-LPS. The control in lane 7 shows that without glutaraldehyde present to effect a covalent crosslinking, the noncovalently crosslinked assemblies of HU-B and f-LPS are dissociated by the effects of SDS and boiling upon HU-B, such that lane 7 is identical to lane 1, since HU-B is seen to be predominantly monomeric due to the presence of SDS (and f-LPS is not stained by Coomassie). These experiments very visually establish that HU-B and f-LPS form large assemblies that become covalently-crosslinked by glutaraldehyde into objects that no longer penetrate the stacking-resolving gel interface of an SDS-PAGE.

#### Fluorescence quenching: f-LPS binding quenches fluorescence in HU-B tryptophan-containing mutants

Figures 3A and 3B show the effects upon tryptophan fluorescence emissions in two tryptophan-containing mutants of HU-B, respectively, F47W HU-B, and F79W HU-B, which fold correctly and retain DNA-binding ability (Supplementary F) upon the addition of micellar f-LPS. The F79W position lies just under the beta hairpin constituting the canonical DNA-binding site in HU-B. The F47W position lies close to the lysine cluster constituting the non-canonical DNA-binding site. The spectra establish that there is less quenching of Trp fluorescence achieved by addition of micellar f-LPS to F79W HU-B than by addition of micellar f-LPS to F47W HU-B. The reasons for this differential response are evident from the differential degrees to which the tryptophan residues are proximal to the DNA-binding sites and exposed to them (see insets in Figures 5A and 5B). This quenching suggests that the binding of micellar f-LPS to HU’s non-canonical DNA-binding site (proximal to F47W) elicits more of a response than binding to the canonical site (proximal to F79W), but that there is a response seen with both mutants, demonstrating yet again that HU-B binds to micellar f-LPS.

#### Changes in birefringence: HU-B binding to immobilized LPS (i-LPS) causes disordering of i-LPS-surfaced liquid crystals

Figure 3 shows effects of binding of HU to liquid crystal-immobilized LPS (i-LPS) upon birefringence of ordered liquid crystals (LCs) of DMOAP (N,N-dimethy-N-octadecyl-3-aminoproyltrimethoxysilyl chloride) coated with 5CB (4-cyano-4’-pentylbiphenyl). In experiments conducted according to protocols established in previous studies,^24^ we show that the 5CB-coated DMOAP liquid crystals exist in an ordered state when exposed to air, in the bulk phase, displaying a dark field under a polarizing microscope, as seen in Figure 3D. When the 5CB-coated DMOAP liquid crystals are placed under water, the water causes a disordering transition in the hydrophobic LC, and a consequent conversion from dark field to bright field, as seen in Figure 3E. However, when micellar f-LPS is immobilized into becoming i-LPS (immobilized f-LPS) upon the LCs of 5CB-coated DMOAP, the LCs once again undergo an ordering transition from bright field to dark field under a polarizing microscope, as shown in Figure 3F, in an air-exposed state. This arrangement shows no further alteration upon being placed under water, as seen in Figure 3G, because the LPS is charged at the end facing the water and its hydrophobic lipid tail interacting with the 5CB is not affected by the water. Thereafter, binding of any reagent to the immobilized i-LPS which can elicit a disordering transition can cause the LCs to go back from being dark field to being bright field even in water, and this is used as a sensor in an assay for protein binding to i-LPS.^24^ Such a disordering transition from dark field to bright field is seen upon the addition of HU-B to liquid crystal-immobilized i-LPS, as seen in Figure 3H, demonstrating that HU-B binds to i-LPS. Figure 3I shows a schematic for the disordering transition caused by protein binding to i-LPS.

### Section 3: Binding of HU to cell-displayed LPS (c-LPS)

#### c-LPS-HU-c-LPS interactions: Flow cytometric evidence of clumping of bacteria through binding of HU-B/HU-A to c-LPS

Figures 4A–4E show flow cytometry data indicating that in the presence of HU-B, as well as HU-A, *E. coli* cells display increased forward scatter (FSC) as well as increased side scatter (SSC) profiles that are diagnostic of the increase in size through cell-clumping. The clumping manifests as a streak on the top-right section of the FSC-SSC scatter plot. Figure 4A shows the FSC-SSC plots for control *E. coli* XL-1Blue cells. When HU-A (Figure 4B) or HU-B (Figure 4C) are added to the *E. coli* cells, there is a tendency for streaks (increased FSC and SSC) to be seen at the top right corner, indicative of clumping of bacteria. When all other conditions are comparable, HU-B appears to cause more clumping than HU-A in these FSC-SSC plots. The distinction between the effect of HU-A and HU-B is clearly also evident in the cell counts plotted against FSC (Figure 4D) and SSC (Figure 4E), i.e., (a) populations seen with addition of HU-A (in green) are only somewhat shifted with respect to the control, unlike the populations seen with addition of HU-B (in blue), and (b) a much more distinctive effect is seen for HU-B that for HU-A in the SSC plots than in the FSC plots.

**Fig. 4.**
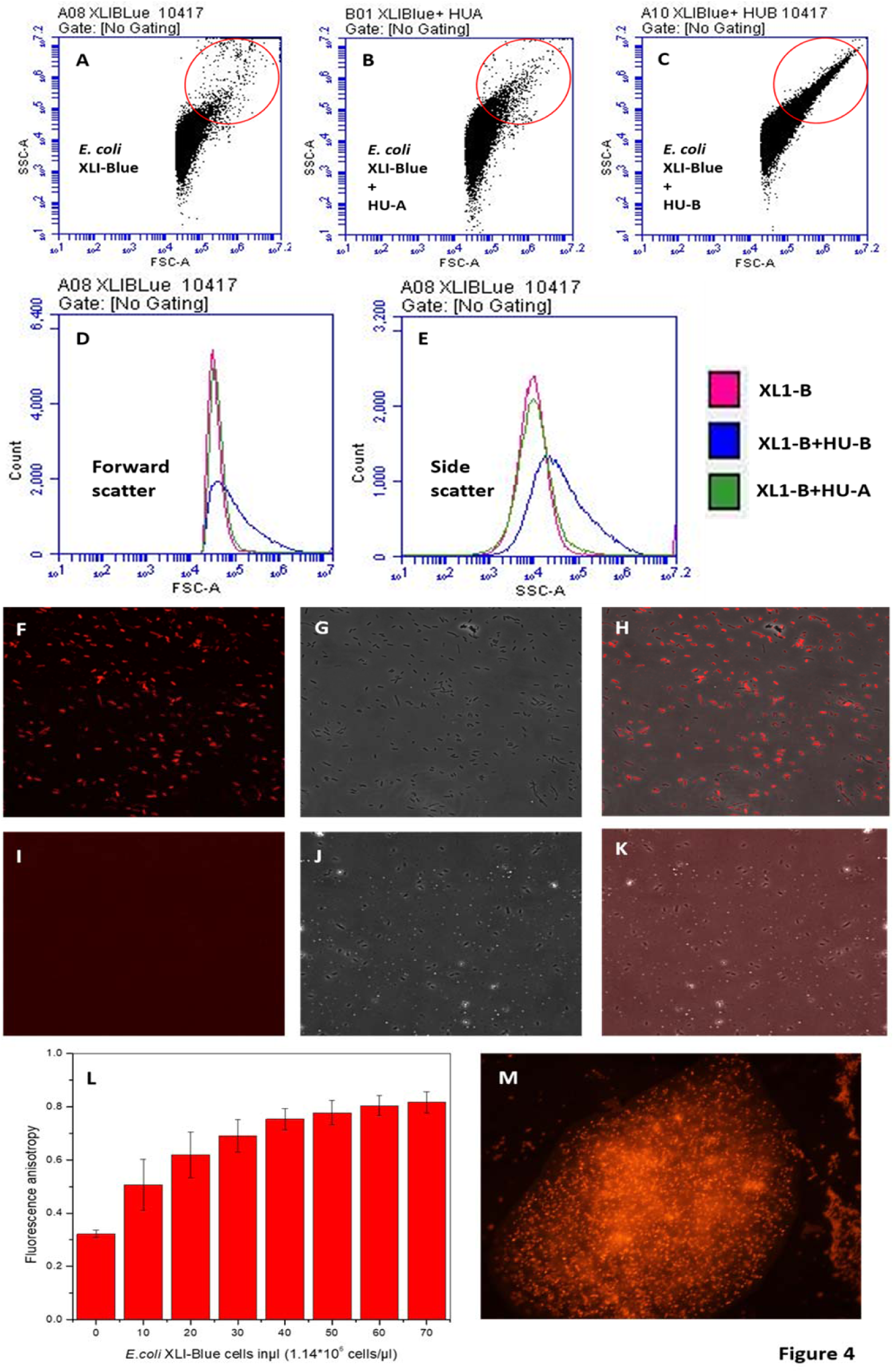
HU-c-LPS and c-LPS-HU-c-LPS interactions. Scatter plots derived from flow cytometry of *E. coli* cells, with monitoring of forward scatter *versus* side scatter using **a**, control XL1-Blue cells, **b**, XLI-Blue cells treated with HU-A, and **c**, XL1-Blue cells treated with HU-B. The ovals (red outlines) represent scatter plot areas with heightened forward and side scatter indicative of bacterial cell clumping. The streak represents clumped/aggregated cells. Distinctions between clumping caused by HU-A and HU-B are observed in **d**, the combined overlay of cell count *versus* forward scatter, and **e**, the combined overlay of cell count *versus* side scatter. **f**, Fluorescence, **g**, phase contrast, and **h**, merged images of XL1-Blue cells treated with exogenously-added Tag-RFP-HU-A. **i**, Fluorescence, **j**, phase contrast, and **k**, merged images of XL1-Blue cells treated with only Tag-RFP. **l**, Changes in florescence anisotropy readout of Tag-RFP-HU-A in the presence of different numbers of XL1-Blue cells. **m**, fluorescence image showing embedment of the bacterial cells in a network of e-DNA created by growing cells without shaking in the presence of HU-B.

The clumping observed is entirely similarly to that observed when poly-D-lysine is added to *E. coli* cells to clump them deliberately (Supplementary Figure 1).

#### HU-c-LPS interactions

Fluorescence microscopic and flow cytometric evidence of binding of RFP-HU-A to bacteria Figures 4F–4K show fluorescence micrographs establishing binding of RFP-HU-A to the surfaces of planktonic bacteria in isotonic buffer of pH 7.4. Control HU-A is non-fluorescent, as it lacks a genetically fused tag-RFP, and so no images are shown for this control. Addition of RFP-HU-A to cells of *E.coli* strain XL-1 Blue elicits localized fluorescence (Figure 4F) which is also co-localized with the bacteria seen in the DIC image (Figure 4G) and in the merged image (Figure 4H). However, addition of RFP alone to *E. coli* cells elicits no localized RFP fluorescence (Figure 4I) corresponding to the DIC image of bacterial cells (Figure 4J), and no overlap of fluorescence and DIC images is seen in the merged image (Figure 4K). This clearly establishes that it is the HU component of the RFP-HU-A fusion which causes RFP-HU-A to co-localize with bacterial cells, and not the RFP domain present in fusion with HU-A in RFP-HU-A. This indicates that HU binds to the surfaces of bacterial cells, since nothing was done to permeabilize cells for this experiment. The indication is that the HU binds to outer membrane LPS (c-LPS), allowing the RFP domain present in fusion to visualize HU titrated to the surfaces of bacterial cells.

It may be noted that the above experiments involved cells that do not overexpress recombinant HU. In separate experiments involving the use of a wide-field fluorescence microscope with capabilities of gathering in-plane images with live bacteria and stacking these into a three-dimensional representation allowing examination of fluorescence within bacteria and outside bacteria, we also observed that cells that overexpress recombinant Venus-HU-B (which sometimes remain syncytial, and form long filaments, and sometimes break into smaller filaments and cells) stick to each other and have a halo of Venus fluorescence outside the cell surface, in addition to that in the cell cytoplasm. (Videos in Supplementary Figure 2).

In flow cytometry experiments, when RFP-HU-A is added to cells instead of HU-A or HU-B alone, the RFP happens to label the cell surface on all cells, and clumps, which display RFP fluorescence (Supplementary Figure 3); however, RFP-HU-A does not cause much clumping, and the streak is much smaller than with HU-B, as noted previously using HU-A. The above observations reconfirm the ability of HU to bind to bacterial cell surfaces through binding to c-LPS. In addition, they suggest that when HU forms large multimeric assemblies, it can cause clumping of bacteria by using different LPS-binding surfaces on such multimers to bind to different bacteria. It is conceivable that when RFP is present in fusion with HU at HU’s N-terminus, this might sterically prevent the formation of tetramers and octamers by dimeric HU polypeptides, even though the presence of the RFP as a domain in the fusion clearly does not interfere with the binding of RFP-HU-A to cells, as was seen in Figures 4F–4K.

#### c-LPS-HU and c-LPS-HU-c-LPS interactions: Fluorescence spectroscopic evidence of increase in anisotropy of RFP-HU-A in the presence of bacteria

Figure 4L shows fluorescence anisotropy data for RFP fluorescence in an experiment in which identical low concentrations of RFP-HU-A are added to different numbers of bacterial cells *(E. coli* strain XL-1 Blue). This results in progressively higher readouts of RFP’s fluorescence anisotropy, indicating that RFP-HU-A is becoming immobilized upon bacterial cells through binding. In this experiment, values of anisotropy higher than 0.4 owe to the high degree of scattering of light which is further increased by the coming together of cells through clumping; however, the transition from a value of 0.2 to higher than a value of 0.4 is clearly indicative of the immobilization of RFP-HU-A on bacterial cells (through binding).

#### c-LPS-HU-c-LPS and c-LPS-HU-DNA-HU-c-LPS interactions: Fluorescence microscopic imaging of bacteria embedded in a matrix of e-DNA

We have already alluded to the generation of e-DNA through cell lysis; ^1^ in particular, through explosive cell lysis during which rod-shaped bacteria lose their shapes and become spheres which then break up to release cytoplasmic contents, and DNA, which is rapidly disseminated in a population of proximally-growing bacterial cells. It is possible that the rapid dissemination of such e-DNA by bacteria occurs through the binding of the e-DNA to bacterial cell surfaces with the assistance of multivalent and abundant nucleoid-associated proteins, including HU. This suggests that addition of HU to a population of growing cells could increase the binding of cells to each other and to e-DNA, promoting greater amounts of explosive cell lysis and generation of even higher amounts of e-DNA. We have found that when bacteria are grown without shaking in the presence, and absence, of exogenously-added HU-B, large bacterial clumps are observed in the petriplates to which HU-B was added. Figure 4M shows that when propidium iodide (PI), a DNA-binding dye, is added to such plates after permeabilizing bacteria, and the clumps are imaged for PI fluorescence, the fluorescing bacteria are observed to be embedded in a matrix of fluorescing e-DNA. This data suggests that HU could play a role in explosive cell lysis by creating physical links between cells, and between cells and DNA, to promote greater lysis through physical stress. It also demonstrates that the availability of additional HU promotes the generation of e-DNA matrices that contain embedded bacteria.

#### f-LPS-HU, DNA-HU and c-LPS-HU interactions: Flow cytometric evidence for inhibition of c-LPS-HU interactions by DNA and f-LPS

Figure 5 shows FSC-SSC scatter plots similar to those shown in Figures 4A–4E, with a dose-dependent reduction of HU-mediated *E. coli* cell clumping observed upon pre-incubation of 4WJ DNA (a 4-way OR 4-stranded cruciform junction) DNA with RFP-HU-A. This suggests that an excess of DNA available for pre-binding to HU saturates its binding sites and reduces scope for use of such sites for binding to c-LPS, upon cell surfaces.

**Fig. 5.**
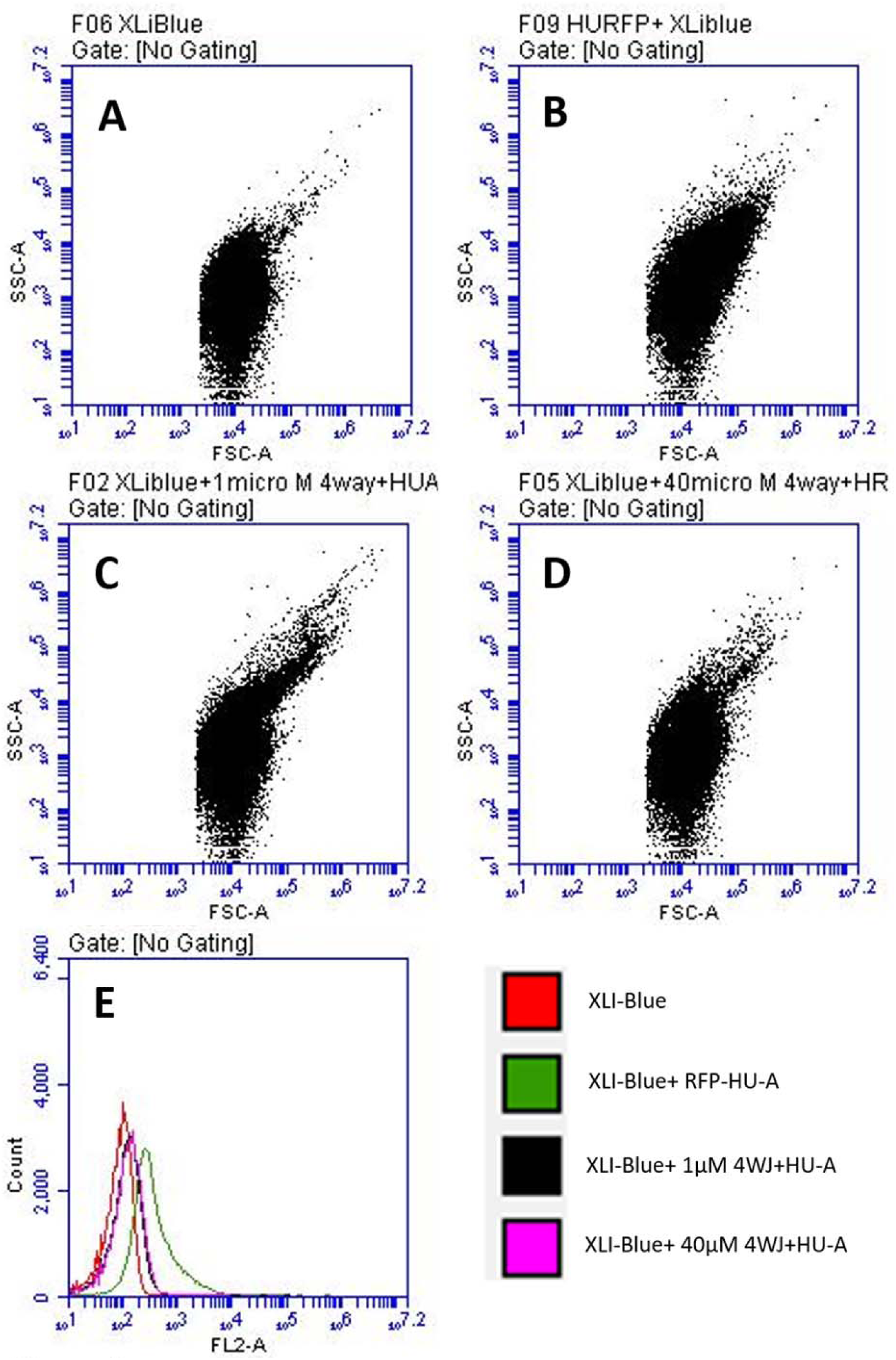
Inhibition of c-LPS-HU-c-LPS interactions (*E. coli* binding and clumping) through pre-incubation of Tag-RFP-HU-A with DNA (4WJ). DNA dose-dependent reduction in intensity of streaks in scatter plots derived from flow cytometry of *E. coli* cells, with monitoring of forward scatter *versus* side scatter using **a**, control XL1-Blue cells, **b**, XL1-Blue cells treated with Tag-RFP-HU-A, **c**, XL1-Blue cells treated with Tag-RFP-HU-A pre-treated with 1 μΜ 4WJ DNA, **d**, XL1-Blue cells treated with Tag-RFP-HU-A pre-treated with 40 μΜ 4WJ DNA. **e**, combined overlay of fluorescence from Tag-RFP-HU-A bound to cells with (and without) pre-treatment with 4WJ DNA; saturation of binding sites on HU-A which are capable of binding to either DNA or c-LPS, by DNA, is observed.

In an analogous set of experiments involving pre-incubation of RFP-HU-A with f-LPS, instead of with 4WJ DNA, Supplementary Figure 4 shows that there is a similar reduction in *E. coli* cell clumping, as the concentration of f-LPS is increased from 0 mg/ml (Supplementary Figure 4A) to 0.5 mg/ml (Supplementary Figure 4B) to 1 mg/ml (Supplementary Figure 4C). This suggests that an excess of f-LPS made available for pre-binding to HU similarly saturates its binding sites and reduces scope for use of such sites for binding to c-LPS. In other words, f-HU molecules that are not pre-bound to something else (e.g., DNA or f-LPS) ae better at causing *E. coli* cell clumping through multivalent binding to c-LPS, than f-HU molecules that are pre-bound to either DNA or f-LPS. It must be borne in mind, of course, that these experiments are merely indicative, and suggestive, of the likely scenario. The actual dissociation constants of f-LPS, c-LPS and 4WJ DNA, for HU-A, and the relative concentrations of c-LPS (dependent upon the number of cells) remain undetermined here. These would, of course, be likely to influence quantitative aspects of the actual data seen.

#### c-LPS-HU interactions: Flow cytometric evidence of inhibition of c-LPS-HU interactions by NaCl

Supplementary Figure 5 shows that increasing concentrations of NaCl are able to screen out HU-LPS interactions sufficiently to abolish the streak that is indicative of *E. coli* cell clumping.

### Section 4: Binding of engineered HU variants to DNA and LPS

#### Characterization of LoodHU-B, LysrHU-B and LoodLysrHU-B through comparison with wild-type HU-B

The wild-type HU and its three variants (described in the first section of this paper) were created and compared with each other, in respect of their ability to fold into dimeric HU. Supplementary Figure 6 shows circular dichroism spectra establishing that HU, and all three variants created to ablate DNA-binding sites, i.e., LoodHU, LysrHU and LoodLysrHU, have comparable structural contents with mean residue ellipticity (MRE) values in the range of −9000 to −9500 deg cm^2^ dmol“^1^ at 208 nm, and negative MRE bands at 208 nm and 222 nm, arising from the helical content of HU. Supplementary Figure 7 shows gel filtration data establishing that HU, LoodHU, LysrHU and LoodLysrHU all have gel filtration elution profiles in which the main population of HU elutes at ∼12 ml, corresponding to a molecular weight of approximately 25 kDa [resulting from the dimerization of two ∼10.6 kDa HU chains, each made up a 1.4 kDa affinity tag (N-MRGSHHHHHHGS) and a 9.2 kDa HU polypeptide, with some disorder in the beta hairpin DNA binding loop adding to the protein’s hydrodynamic volume].

#### 4WJ DNA binds to HU-B, LoodHU-B and LysrHU-B but not to LoodLysrHU-B

Above, we have presented data supporting the folding and dimerization of (i) HU-B (i.e., wild-type HU-B), (ii) LoodHU-B (i.e., loop-deleted HU-B lacking the canonical DNA-binding site, in which the DNA-binding beta hairpin loop has been removed and replaced by a glycine-serine linker in each monomer), (iii) LysrHU-B (i.e., lysine cluster-replaced HU-B lacking the non-canonical DNA-binding site, in which mutations K83A and K86A have been introduced together; note, the K83A and K86A mutations were also introduced individually, but these variants have not given special names, although data regarding these is also shown in this section) and (iv) LoodLysrHU-B (i.e., loop-deleted and lysine cluster-replaced HU-B lacking both canonical and non-canonical DNA-binding sites). Similarly, some of these DNA binding site-ablated molecular species were also created using HU-A. Once we were satisfied that the DNA-binding site-ablated variants were folded and dimeric like HU-B/HU-A, we examined all available molecular species in respect of their ability to bind to DNA. We found that HU-B, LoodHU-B, and LysrHU-B (Figures 6A and 6B) as well as HU-A, LoodHU-A and LysrHU-A (Figure 6C) are capable of binding to 4WJ DNA, as judged by their ability to elicit an electrophoretic mobility shift. However, the LoodLysrHU-B and LoodLysrHU-A variants lacking both canonical and non-canonical DNA binding sites showed no ability to bind to DNA. In Figures 6A–6C, control DNA (4WJ) is shown in lanes 1, 7 and 11. HU-B, LoodHU-B and HU-B containing individual K83A and K86A mutations are all seen to bind 4WJ DNA (lanes 2-5) as is LysrHU-B (lane 6). However, LoodHU-B additionally lacking either of the K83 or K86 lysine residues, or both lysine residues (i.e., partial and total LoodLysrHU-B variants) fail to bind to 4WJ DNA (lanes 8-10). Similarly, with HU-A and its LoodHU-A and LysrHU-A variants, it is seen that HU-A lacking K83, K86 or both lysine residues (i.e., partial or total LysrHU-A) is still able to bind to 4WJ DNA, as would be expected, because of the retention of the canonical DNA-binding site. Interestingly, exactly as seen with HU-B, when the canonical DNA-binding site is absent (i.e., LoodHU-A) and some or all of the non-canonical site is also absent, i.e., when the variant additionally lacks either K83 or K86 or both lysine residues (i.e., partial or total LoodLysrHU-A), no binding of 4WJ DNA is observed. From all of this data is it incontrovertibly clear that at least one of the two DNA-binding sites (the canonical loop, or the non-canonical double-lysine cluster) is required by HU for binding of DNA. Compromise of either site is tolerated and the HU molecule still binds to DNA, but compromise of both sites through loop deletion, or removal of one or both lysine residues, is not tolerated, and there is no DNA binding by such variants. In the section below, we present evidence based on glutaraldehyde crosslinking experiments that ablation of the same two sites also completely abrogates LPS binding, whereas individual ablation of the sites does not elicit the same effect.

**Fig. 6.**
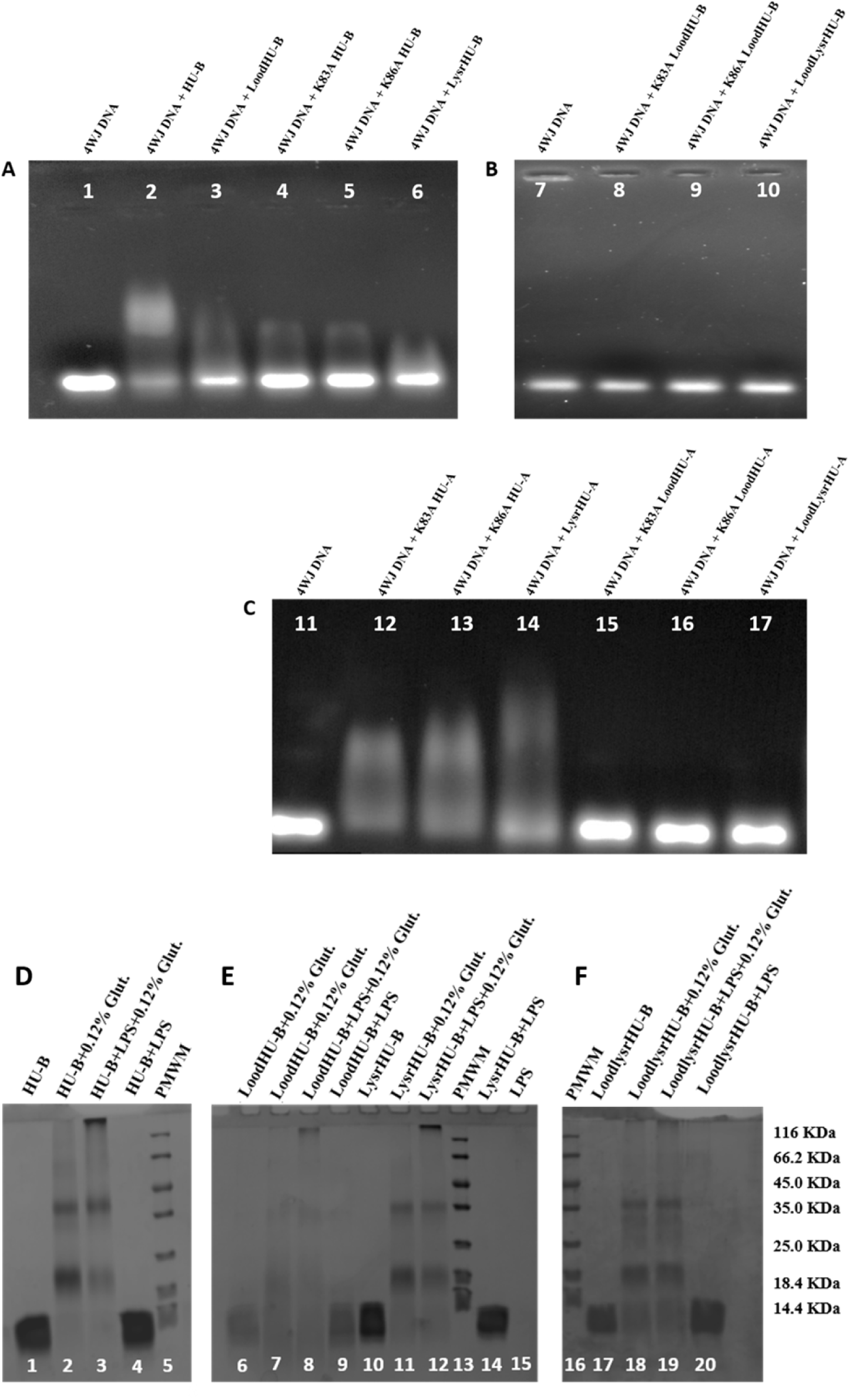
Binding of 4WJ DNA and f-LPS by HU-B, HU-A and their Lood, Lysr and LoodLysr variants. Electrophoretic mobility shift assay (EMSA) performed on agarose gels with EtBr-stained 4-way junction (4WJ) DNA bound to various HU variants as shown in **a**, for individual HU-B variants with ablated canonical site, *or* partially/totally ablated non-canonical site. **b**, for HU-B variants with ablated canonical site *and* partially/totally ablated non-canonical site, or **c**, for individual HU-A variants with ablated canonical site *or* partially/totally ablated non-canonical site, *or* HU-A variants with ablated canonical site *and* partially/totally ablated non-canonical site. SDS-PAGE gel assays for glutaraldehyde-based covalent crosslinking of HU-B into HU-B multimers (dimers, tetramers and hexamers) and HU-B-f-LPS high molecular weight (HMW) forms unable to cross the stacking-resolving gel interface, as shown in **d**, for HU-B. **e**, for individual HU-B variants with ablated canonical site *or* totally ablated non-canonical site, **f**, for HU-B variants with ablated canonical site *and* totally ablated non-canonical site.

#### f-LPS binds to HU-B, LoodHU-B and LysrHU-B but not to LoodLysrHU-B

We next explored the binding of f-LPS by HU-B and its three variants, LoodHU-B, LysrHU-B, and LoodLysrHU-B, using the glutaraldehyde crosslinking experiment (as shown earlier in Figure 2H), to establish whether glutaraldehyde crosslinks LPS, on the one hand, and HU-B and its above variants, on the other hand into large covalently crosslinked assemblies that fail to enter the resolving SDS-PAGE gel, remaining at the interface of the stacking and resolving gels. In Figures 6D–6F, lanes 5, 13 and 16 show the electrophoretic migration of protein molecular weight markers of known size (mentioned at the right end of the figure). Lanes 1, 6, 10 and 17, show the electrophoretic migration of HU-B, LoodHU-B, LysrHU-B and LoodLysrHU-B, below the 14.4 kDa protein molecular weight marker. Lanes 2, 7, 11 and 18, show the crosslinking by glutaraldehyde of HU-B and all three variants of HU-B into dimers (just above the 18.4 kDa protein molecular weight marker), tetramers (just above the 35 kDa protein molecular weight marker) and, in some lanes, hexamers (around the 66.6 kDa bands). Lane 3 shows that glutaraldehyde crosslinks HU-B and f-LPS into aggregates that cannot enter the resolving gel, and lane 4 shows that when no glutaraldehyde is present there is no crosslinking. Entirely similar results with LoodHU-B are seen in lanes 8 and 9, and with LysrHU-B in lanes 12 and 14. However, with LoodLysrHu-B, the lanes with and without glutaradehyde are nearly identical, displaying crosslinking of LoodLysrHU-B into dimers, and tetramers, but not into the aggregates that are unable to enter the resolving gel, at the resolving-stacking gel interface.

## Concluding Discussion

The *E. coli* nucleoid-associated proteins (NAPs), HU-A, and HU-B, share ∼69 % amino acid sequence identity. We have shown that both HU isoforms are capable of binding to free lipopolysachharide (f-LPS), as well as to cell-displayed LPS (c-LPS) on the outer membranes of bacteria. HU possesses two types of DNA-binding sites. The first of these sites was discovered along with the determination of the structure of HU in complex with DNA. The other site was discovered subsequently, based on small angle X-ray scattering studies. We refer to these DNA-binding sites are HU’s ‘canonical’, and ‘non-canonical’, DNA-binding sites, respectively. Through bioinformatics-based distance measurements we have shown that certain pairs of positively charged amino acid residues at each of these sites (i.e., residues R58 and R61 at the canonical DNA-binding site, and residues K83 and K86 at the non-canonical DNA-binding site) happen to be perfectly positioned for binding of the phosphate moieties present in the lipid-A head groups of LPS. Further, using microscale thermophoresis, difference absorption spectroscopy, dynamic light scattering, biolayer interferometry, glutaraldehyde crosslinking and polarized microscope-based birefringence studies using liquid crystals that HU and LPS interact. Using microscale thermophoresis we measured an affinity in the range of a few tens of micromolar (∼34 μM); however, we do not lay much store by the accuracy of this particular affinity measurement, because we are not confident about the homogeneity of the commercially-sourced LPS, and unsure about the fraction of the LPS that is present in micellar form, since most of our experiments (except for the light scattering experiments) were conducted using LPS concentrations in the post-CMC (critical micellar concentration) range of LPS concentrations.

Towards the end of the paper, we have also presented evidence from experiments in which we genetically ablated the canonical and non-canonical DNA-binding sites of HU, both individually and together, to demonstrate that each site is capable of binding to DNA, as well as to LPS, and that genetic variants lacking both types of sites are unable to bind to either DNA or LPS. This establishes, in our view, that there is a complete physical coincidence of the DNA-binding and LPS-binding sites of HU. In hindsight, this is not altogether surprising, considering that the lipid-A headgroup of LPS contains an arrangement of sugars in conjunction with phosphate groups (4-phosphoβ-GlcN-(1,6)-α-GlcN-1-phosphate), somewhat akin to the presence of sugars and phosphates in the backbone of DNA. In DNA, the phosphate groups are constituent parts of phosphodiester bonds, whereas in LPS they are terminal phosphates. Thus, each phosphate group in DNA carries a single negative charge, whereas each of the two phosphate groups in LPS carries two negative charges. Further, each sugar in DNA is a pentose (ribose) sugar, whereas each sugar in the lipid A headgroup of LPS is a hexose (glucosamine, or GlcN) sugar. Further, the two GlcN sugars in LPS have no intervening phosphate group and so the headgroup of LPS is only notionally, or nominally, like DNA, in that it is capable of presenting phosphate groups to the DNA-binding site of a DNA-binding protein, in conjunction with arrangements of carbon, hydrogen and oxygen atoms derived from a sugar. The important thing is that LPS can bind to the DNA-binding site of a DNA-binding protein by presenting sugar-phosphate moieties to the site.

Our further studies with *E. coli* cells indicate that HU is capable of using its ability to bind to LPS to become visibly titrated onto the surfaces of *E. coli* cells (with consequent increase in the anisotropy of fluorescence of RFP domains present in fusion with HU), and to cause the clumping of cells detectable by flow cytometry, by acting as a glue that allows negatively-charged cell surfaces (which could otherwise be expected to repel each other) to bind to positively-charged multimers of HU. We have shown that *E. coli* cells expressing HU tend to attach to each other after division, suggesting the involvement of some proteins on the cell surface (which could be HU). We have also shown that fusions of YFP/Venus and HU appear outside the cell, in association with the cell surface. In fact, it was this observation which first caused us to wonder about the presence of HU outside the cell, and about its possible association with the cell surface.

There is evidence that HU can appear outside the cell in the extracellular material that constitutes biofilms.^5^ There is also evidence that HU can be secreted by type IV pathways,^11^ or simply thrown out of cells through explosive cell lysis.^9^ We propose that HU present in the extracellular medium in a developing biofilm binds to the surfaces of adjoining cells that are also in motion, and that this results in uneven stresses that become the cause for explosive cell lysis. In fact, videos of such lysis indicate that the expelled DNA (which turns into the e-DNA matrix of a biofilm) is bound by cells and dragged-around within a colony of cells.^9^ We feel that this is because the expelled DNA is decorated with molecules of HU, and that not all of the DNA-binding sites on such HU molecules are engaged in the binding of DNA, with some remaining vacant and available for interactions with the surfaces of bacterial cells. It is possible that vacant DNA-binding sites bind to c-LPS on cell surfaces, allowing cells to bind to e-DNA through interactions with DNA-bound HU. In fact, this has the potential for becoming a self-perpetuating process, such that each event of explosive cell lysis could become the cause of the next event of explosive cell lysis, simply by making more e-DNA available for cells to bind to (and feel physically stressed by), as they grow, divide and move around inside a bacterial colony. Indeed, we have shown evidence that exogenous addition of HU to bacterial cells in non-shaken cultures leads to the generation of large propidium iodide-binding, DNA-rich entities (i.e., simulacrums of biofilms) in which bacterial cells are found to be embedded. This suggests that addition of HU can begin the process of biofilm formation even with freshly growing and dividing bacteria which are most likely to possess sites of weakness in their outer cell membranes and cell walls, through the promotion of more e-DNA expulsion. Once again, there is evidence in the literature to suggest that addition of HU reinforces the formation of biofilms.^5^

Thus, we emphasize that HU could be a central player in biofilms by being present in great abundance, by being present everywhere upon the e-DNA matrix within biofilms, and also by being able to bring together e-DNA and negatively-charged surfaces of bacterial cells, as well as clump cells, as shown in this paper. Therefore, we emphasize that the LPS-binding and cell-clumping abilities of HU, far from being interesting curiosities, could actually be central factors in the mechanism of association of *E. coli* cells to form biofilms, especially under conditions of exhaustion of nutrients and/or starvation, since under such conditions, the death of a few cells (and the HU-bound DNA expelled thereby) could rapidly scale up cell-cell associations, stresses upon cells, instances of DNA expulsion, participation of cells in clumping and in binding to DNA-bound HU. LPS-binding by HU could thus cause cells to stick to each other, as we have shown, as well as cause cells to stick to bits of e-DNA (themselves networked through binding to multimeric forms of HU), to form the foundations of biofilms. Of course, it is reasonable to assume there would be likely constant competition between e-DNA, and LPS, for binding to HU’s DNA-binding sites. We have shown (i) that DNA-bound HU binds less to c-LPS (to cause less clumping), (ii) that f-LPS-bound HU also binds less to c-LPS (to cause less clumping), and (iii) that salt interferes electrostatically with the interactions of HU with c-LPS (to cause less clumping). It is already known that antibodies raised against the DNA-binding tips (i.e, the canonical beta hairpins) of HU cause dislodgement of bacteria from biofilms.^6^ All of the available evidence, therefore, points towards a very key involvement of HU in the formation and stabilization of biofilms. It may be noted that all bacteria have negatively-charged surfaces, and that if they do not possess c-LPS on their surfaces, they possess another negatively charged sugar-phosphate arrangement involving a different molecule lipotechoic acid.

### Methods

#### Bacterial strains, media, plasmids and protein expression

The XL-1 Blue strain of *E.coli* K-12 was used for all experiments. Cells were grown using LB medium. Expression of all HU-based proteins (including engineered variants of HU) was carried out using these cells, for which cells were transformed by pQE-30 (Qiagen) expression vectors incorporating genes encoding proteins of interest inserted in the vector’s multiple cloning site. Proteins were purified using standard Ni-NTA affinity purification IMAC methods (Qiagen Expressionist) under non-denaturing conditions. Attached genomic nucleic acids fragments and bound proteins were removed from HU during purification through washing of all Ni-NTA-bound HU protein forms (wild type and mutants/variants) with 15 column volumes of 2 M NaCl, leading to dissociation and removal of all contaminant DNA and associated proteins.

#### Recombinant (engineered) proteins and other reagents

Genes encoding HU-A and HU-B were PCR amplified from *E. coli* K-12 genomic DNA and cloned into the multiple cloning site of the pQE-30 vector between the Bam HI and Hind III restriction enzyme sites, such that the two proteins (and all of their mutants, deletion or truncation variants, and fluorescent protein fusions, as described below) could be produced with an N-terminal 6xHis affinity tag facilitating purification: (i) HU-A wild type; (ii) HU-B wild type; (iii) RFP-HU-A, i.e., HU-A with red fluorescent protein (tagRFP) present in fusion at HU’s N-terminus, without any linker; (iv) Venus-HU-B, i.e., HU-B with yellow fluorescence protein/Venus present in fusion at HU’s N-terminus, without any linker; (v) tryptophan-containing HU-B mutant, F47W HU-B; (vi) tryptophan-containing HU-B mutant, F79W HU-B; (vii) LoodHU-B, i.e., HU-B in which residues 52-74 (beta hairpin loop containing HU’s canonical DNA-binding site residues) are ablated/deleted and replaced by an 11 amino acids-long glycine-serine linker (N-SGGGGSGGGGS-C); (viii) LysrHU-B, i.e., HU-B in which residue mutations K83A and K86A (at HU’s non-canonical DNA-binding double-lysine cluster) effect lysine residue by alanine residues; and (ix) LoodLysrHU-B, i.e., HU-B combining the LoodHU-B and LysrHU-B mutations.

All mutants and variants were verified through DNA sequencing. Oligonucleotides used for PCR and splicing by overlap extension (SOE) PCR-based mutagenesis or protein fusions were sourced from Integrated DNA Technologies (IDT), or Sigma. Lipopolysaccharide (LPS) was sourced from Sigma (Catalog No. L-2630-25MG). Four-way junction (4WJ) DNA was sourced in the form of four independent oligonucleotides from IDT or Sigma through contract synthesis, and assembled into 4WJ DNA through addition of the following four oligonucleotide strands to each other in equimolar amounts (Strand 1: 5’-CCCTATAACCCCTGCATTGAATTCCTGTCTGATAA-3’; Strand 2: 5’-GTAGTCGTGATAGGTGCAGGGGTTATAGGG-3’; Strand 3: 5’-AACAGTAGCTCTTAATTCGAGCTCGCGCCCTATCACGA CTA-3’; Strand 4: 5’-TTTATCAGACTGGAATTCAAGCGCGAGCTCGAATAAGAGCTACTGT-3’). Restriction enzymes used for recombinant DNA work were sourced from Thermo Fisher Scientific. DNA polymerase and ligase enzymes were sourced from New England Biolabs. All other media, chemicals and reagents, including poly-D-lysine (Catalog No. P6407-5MG) were sourced from Hi-Media or Sigma, or from individual manufacturers of instruments for consumables associated with specific instruments.

#### Instrument-based spectroscopic, microscopic, cytometric and other analyses

*UV-visible absorption spectral measurements* spectra for protein concentration estimation were collected on a Varian 50-Bio spectrophotometer, using a micro-cuvette with a path length of 0.3 cm for standard measurements. For *DAS (difference absorption spectroscopy)* measurements, a tandem quartz cuvette of 1 cm path length was used, incorporating two tandem compartments of 0.45 cm path length each. The two tandem compartments were filled with 1 ml of 2 mg/ml LPS, and 1 ml of 10 μM RFP-HU-A, respectively. Mixing of the contents was done through inversion of the cuvette after collection of the baseline. *Fluorescence spectral measurements* for collection of *fluorescence emission* spectra as well as *fluorescence quenching* upon LPS binding to tryptophan-containing variants of HU-B, were made using 295 nm excitation and 5 nm excitation and emission slid-widths on a Varian Eclipse spectrofluorimeter, using a 0.3 x 0.3 cm quartz cuvette, and HU-B (F47W or F79W) protein of 0.65 mg/ml concentration and LPS concentrations varying from 0.14-0.56 mg/ml. *Fluorescence anisotropy* measurements were made for red fluorescent protein (RFP) on a Horiba Fluoromax spectrofluorimeter, using 550 nm excitation, and excitation and emission slit-widths of 2.5 nm each, using 10 nM RFP-HU-A and varying cell numbers. *Circular dichroism* spectra were collected to estimate protein secondary structural content using a BioLogic M0S-500 spectrometer, using a cuvette of 0.1 cm path length, and protein concentrations in the range of 0.25-0.35 mg/ml. *Dynamic light scattering* measurements of protein size were made using Wyatt Dawn 8+ instrument and ASTRA software, using a protein concentration of 0.5 mg/ml and an LPS concentration of 2 mg/ml. *Biolayer interferometry* measurements were made using a ForteBio BLItz instrument and Ni-NTA-derivatized tips from ForteBio to examine interactions between tip-bound HU and LPS, using an HU-B concentration of 6 μM and LPS of 1 mg/ml concentration, and PBS buffer of pH 7.4. *Microscale thermophoresis* measurements were made on a Nanotemper Monolith NT-115 instrument with 16 capillaries, with fluorescence excitation through a 550 nm laser to examine protein diffusion as a function of LPS to estimate HU-LPS binding. The RFP-HU-A protein concentration used for 250 nM, and the LPS concentration was 2.5 mg/ml in the first capillary and serially-diluted through halving of concentrations over the remaining 15 capillaries. The temperature jump involved heating by 2 °C. *Flow cytometry* measurements of bacterial clump sizes and the fluorescence associated with binding of RFP-HU-A to bacteria were made on a Becton-Dickinson Accuri C-6 instrument. *Gel filtration chromatography* measurements of protein elution volume and molecular weight for HU-B and its variants were made using Superdex-75 10/300 GL columns on an Akta Purifier-10 GE Healthcare instrument. *Fluorescence microscopy* images were collected to examine propidium-iodide labelled permeabilized *E. coli* cells embedded in a DNA matrix using a Nikon Eclipse Ti-u microscope. Confocal *fluorescence microscopy* images were acquired using a table-top Olympus FLUOVIEW FV10i microscope to examine RFP-HU association with XL-1Blue *E. coli. Wide-field fluorescence* and DIC *deconvolution microscopy* images and videos were collected using a wide-field, high-resolution, Delta Vision Deconvolution microscope [Model DV Elite, GE Healthcare] equipped with solid state illumination and a 1.4 megapixel monochrome CCD camera [CoolSnap HQ2, Photometrics]. *Analytical electrophoresis* was performed using a Bio-Rad Mini-Protean Tetra vertical electrophoresis setup for glutaraldehyde-based crosslinking studies of LPS with HU-B (and variants) using 15% SDS-PAGE gels and Coommassie Blue G-250 protein staining according to standard methods, and using a Bio-Rad Wide Mini-Sub Cell GT submarine set up for electrophoretic mobility shift assays involving 4WJ DNA and HU-B (and variants) using 0.7 % or 1 % agarose gels and ethidium bromide DNA staining. *Protein structural and distance analyses/representations* used PYMOL software from Schrodinger.

## Data availability statement

All data generated or analysed during this study are included in this published article (and its supplementary information files). However, raw data files generated by instruments during and/or analysed during the current study (are available from the corresponding author on reasonable request.

## Competing Interests

The authors declare that there are no competing interests.

## Author Contribution

Bhishem Thakur, Kanika Arora and Archit Gupta performed experiments in decreasing order of contribution. Bhishem Thakur and Purnananda Guptasarma planned and designed experiments and analysed data. Bhishem Thakur and Purnananda Guptasarma wrote the paper. Purnananda Guptasarma supervised the work.

## Acknowledgements

We thank Ms. Ipsita Pani and Dr. Santanu Pal of the Department of Chemical Sciences, IISER Mohali, for making their liquid crystal-based assay for protein-LPS binding available for us to test whether HU binds to LPS using their assay. We thank Ms. Angel for technical assistance. PG thanks the Ministry of Human Resource Development, Government of India, for the grant (MHRD-14-0064) of a Centre of Excellence in Frontier Areas of Science and Technology in the area of Protein Science, Design and Engineering. BT, KA, and AG, respectively, thank the Council of Scientific & Industrial Research, the University Grants Commission and the Department of Biotechnology for doctoral research fellowships.

---

**Supplementary Fig. 1.**
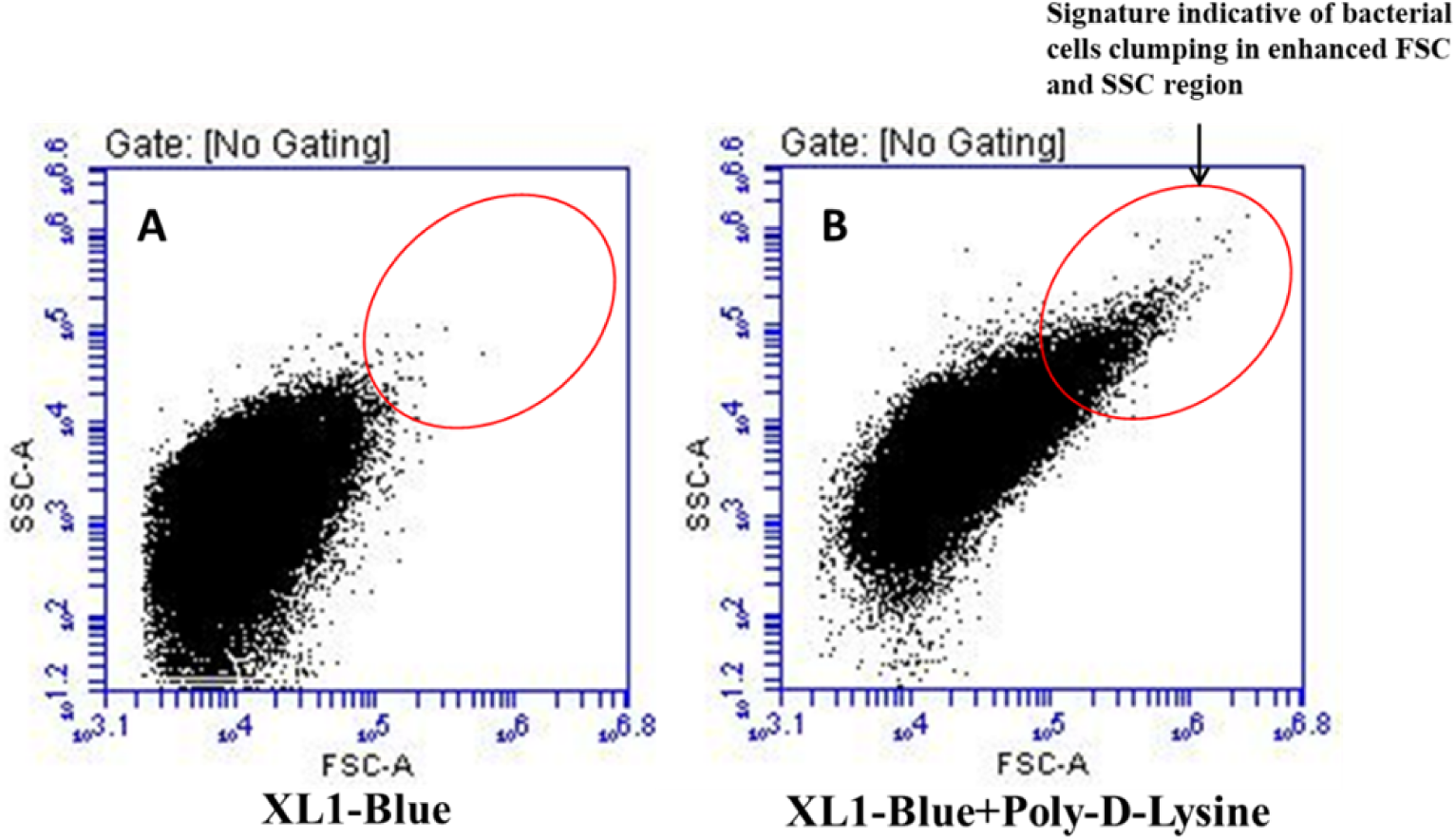
Poly-D-lysine mediated bacterial cells clumping. **a**, Scatter plot of for the untreated XL1-Blue cells plotted between FCS-A, SSC-A, X and Y axis respectively. **b**, Scatted plot of for the XL1-Blue cells after treating with poly-D-lysine.

**Supplementary Fig. 2.**
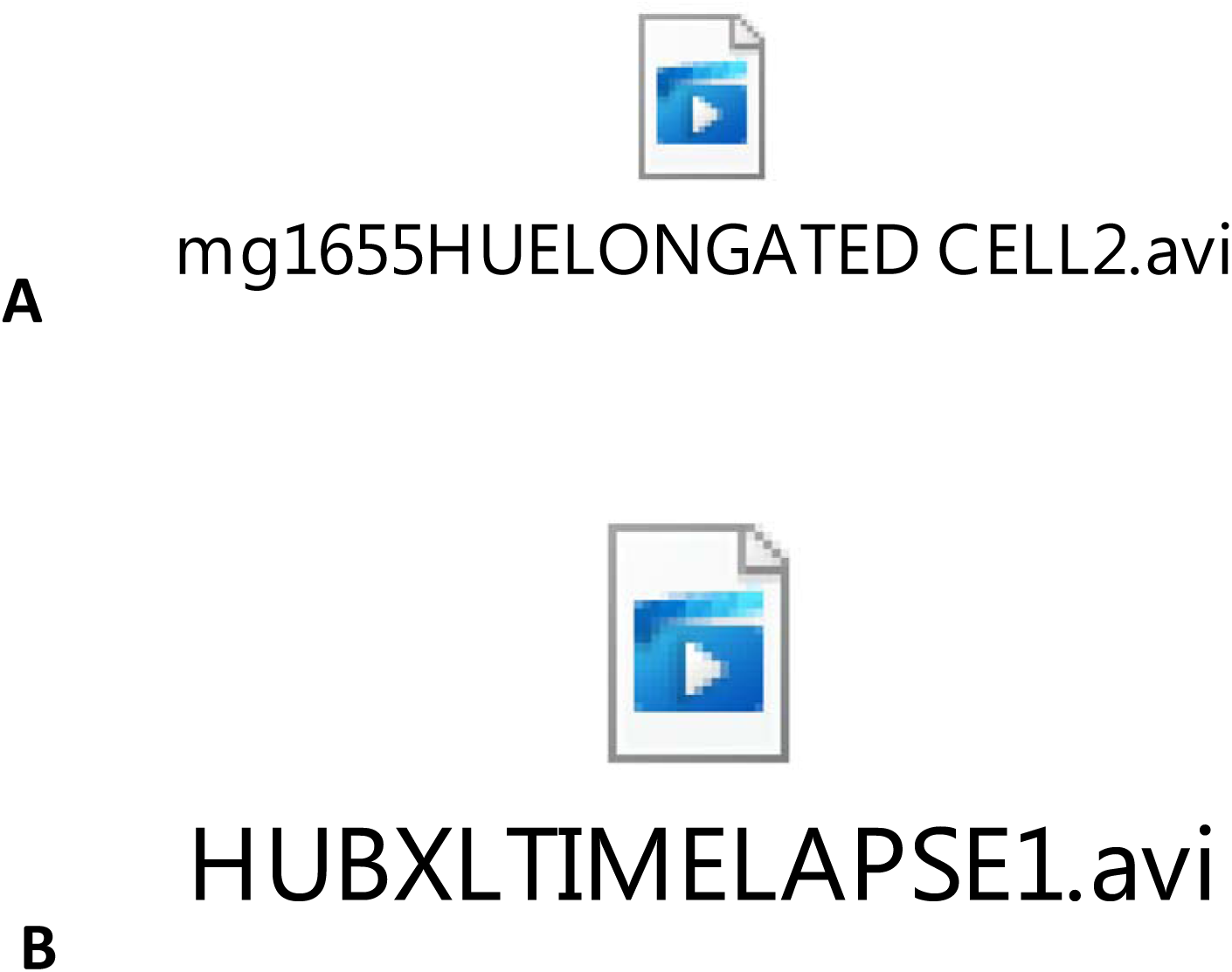
Presence of Venus-HU-B on the surface of expressing *E. coli* MG1655 cells. **a**, Click on the above Windows Media Player video tab to play a video of an elongated (non-separated) syncytial *E.coli* (strain MG1655) cell expressing Venus-HU, in which the progressive viewing of different z-axial depths reveals that there is a halo of Venus-HU-B around the surface of the cell which is distinct from the Venus-HU-B present inside the cell in association with the cell’s chromosomal nucloied. The cell is on agar. The sections of the halo around the cell which are above the agar are much more distinctly visible than the sections in association with the agar. **b**, syncytial cell expressing Venus-HU-B breaking to generate smaller filament and individual cells displaying a tendency to stick to each other along their surfaces.

**Supplementary Fig. 3.**
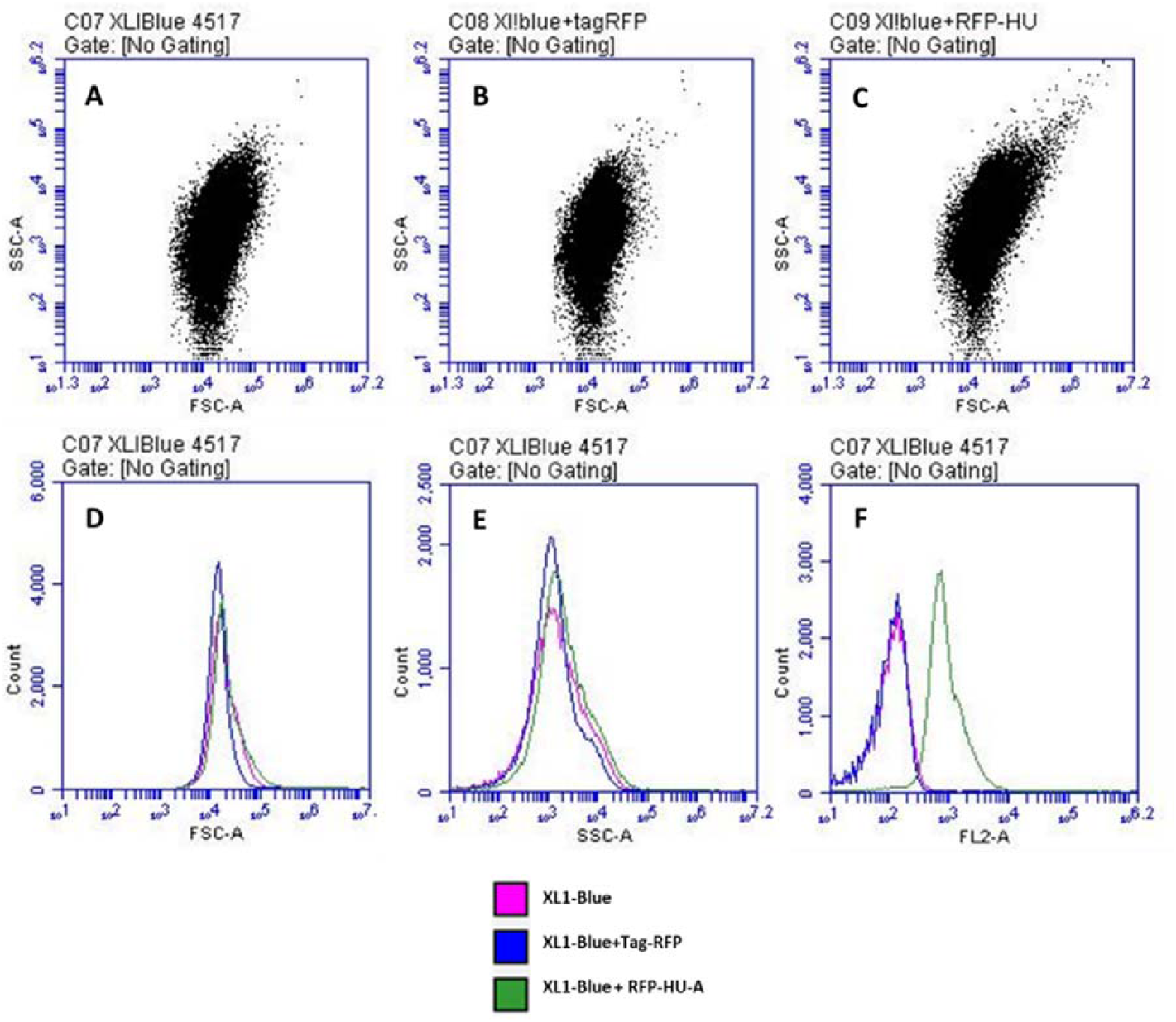
Tag-RFP-HU-A binds to the c-LPS without causing much clumping proved by flow cytometry. **a, b, c**, Scatter plot of for the untreated XL1-Blue cells and the cells incubated with Tag-RFP and Tag-RFP-HU-A proteins. **d, e**, Overlay of the histogram for three sets of bacterial cell counts plotted with FSC-A and SSC-A values. **f**, Overlay of the histograms of three sets of the cells to compare the amount of the fluorescence associated with them.

**Supplementary Fig. 4.**
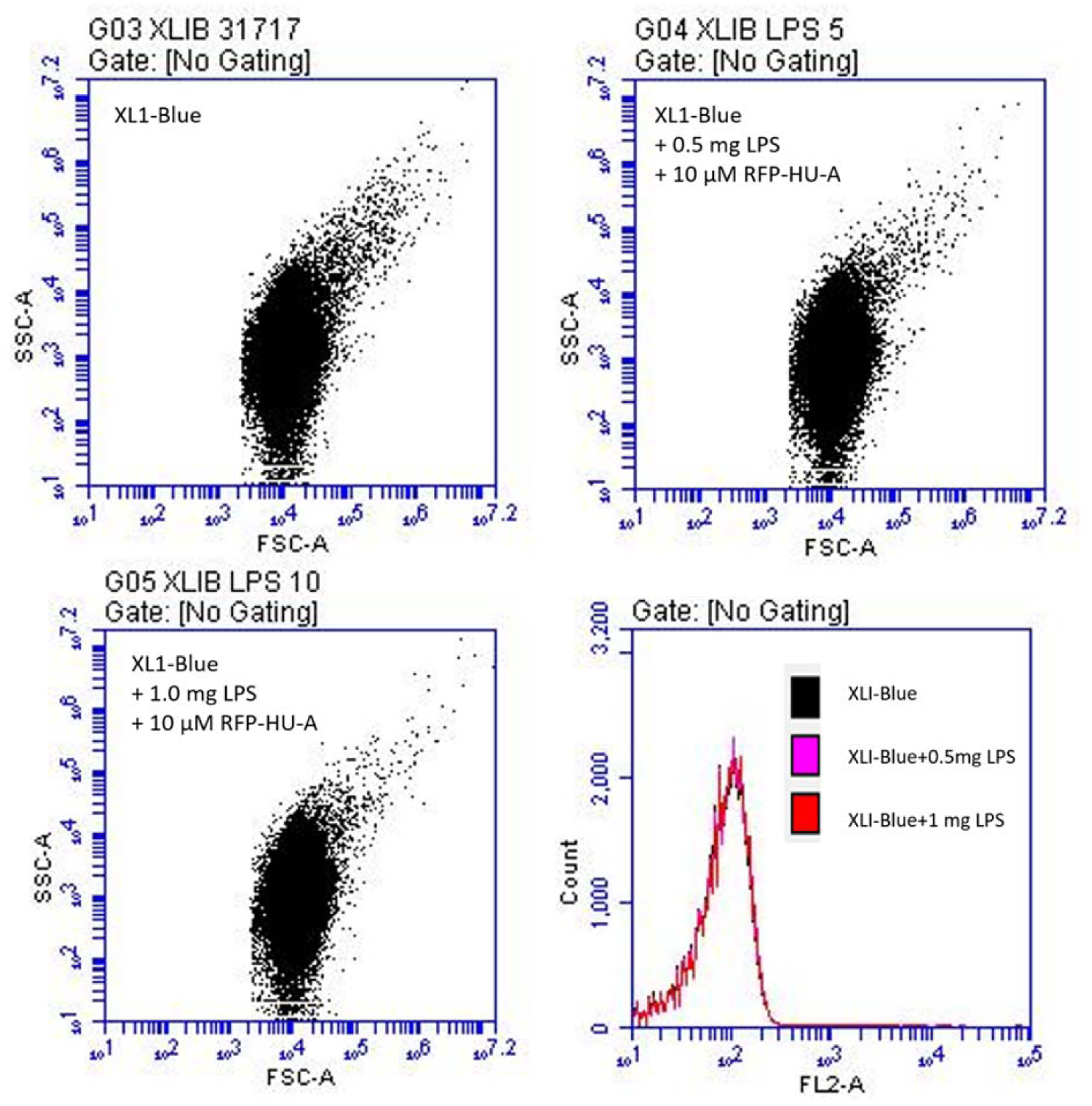
Inhibition of c-LPS-HU-c-LPS interactions *(E. coli* clumping) through pre-incubation of RFP-HU-A with f-LPS. f-LPS dose-dependent reduction in intensity of streaks in scatter plots derived from flow cytometry of *E. coli* cells, with monitoring of forward scatter *versus* side scatter using **a**, control XL1-Blue cells, **b**, XL1-Blue cells treated with 10 μM RFP-HU-A pre-treated with 0.5 mg/ml f-LPS, **c**, XL1-Blue cells treated with 10 μM RFP-HU-A pre-treated 1.0 mg/ml f-LPS. **d**, combined overlay for all cells with (and without) pre-treatment of RFP-HU-A with f-LPS; saturation of binding sites on HU-A by f-LPS lead to lack of RFP-HU-A binding to cells.

**Supplementary Fig. 5.**
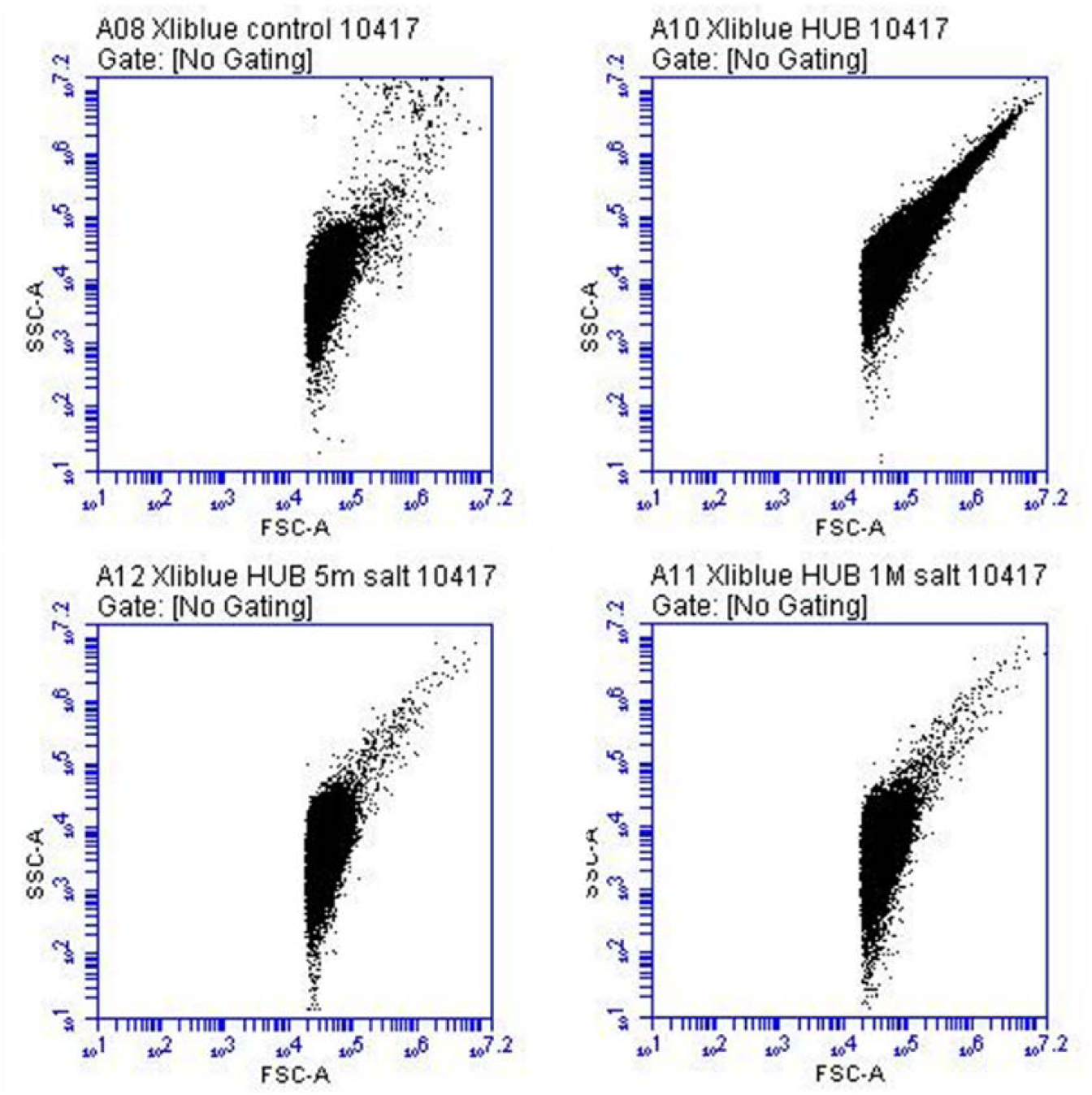
Inhibition of c-LPS-HU-c-LPS interactions (*E. coli* clumping) through pre-incubation of HU-B with NaCl. NaCl dose-dependent reduction in intensity of streaks in scatter plots derived from flow cytometry of *E. coli* cells, with monitoring of forward scatter *versus* side scatter using **a**, control XL1-Blue cells, **b**, XL1-Blue cells treated with HU-B, **c**, XL1-Blue cells treated with HU-B in presence of 0.5 M NaCl. **d**, XL1-Blue cells treated with HU-B in presence of 1.0 M NaCl.

**Supplementary Fig. 6.**
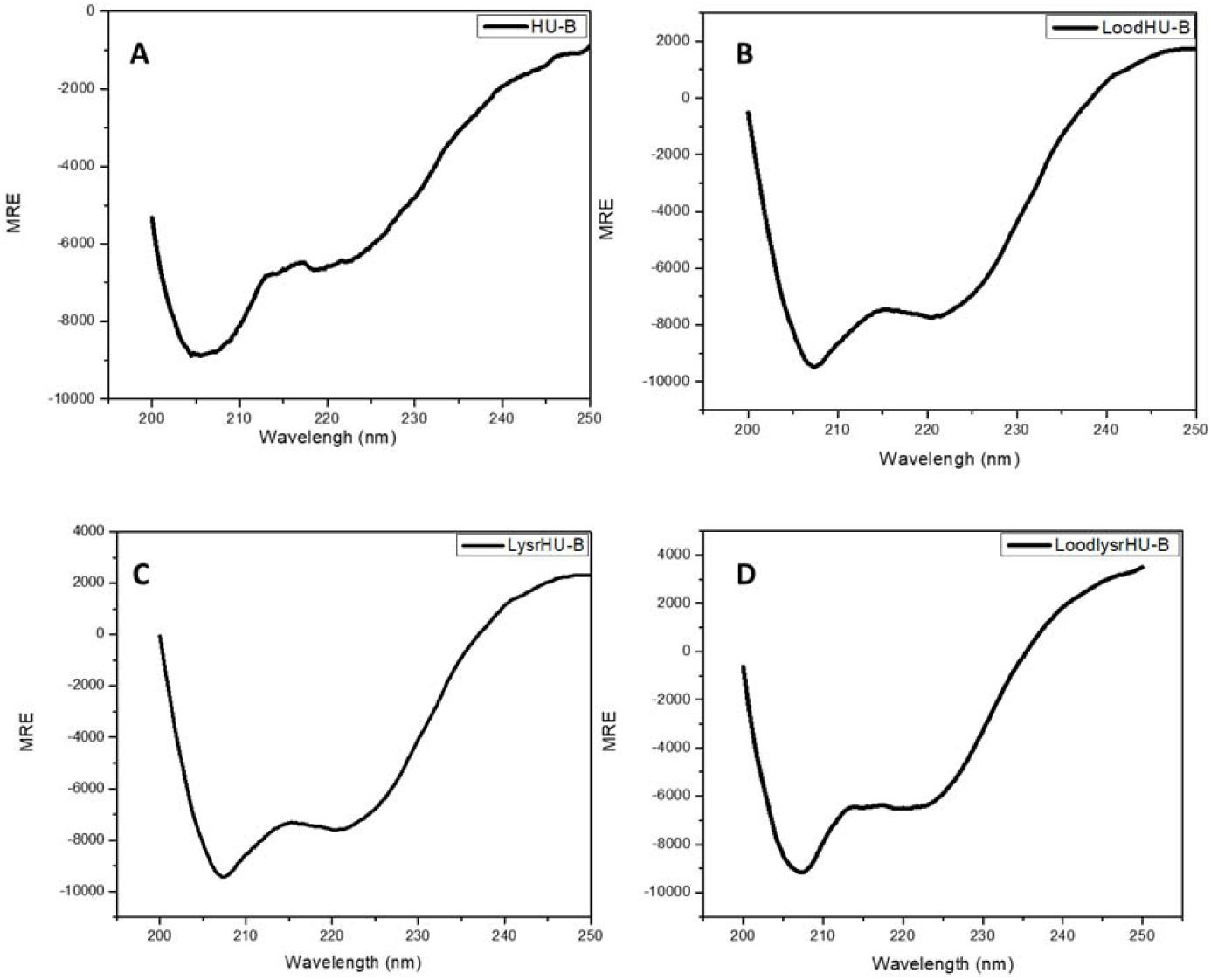
Circular Dichroism (CD) spectra of HU and its DNA-binding site-ablated variants. **a**, HU-B. **b**, LoodHU-B. **c**, LysrHU-B. **d**, LoodLysrHU-B.

**Supplementary Fig. 7.**
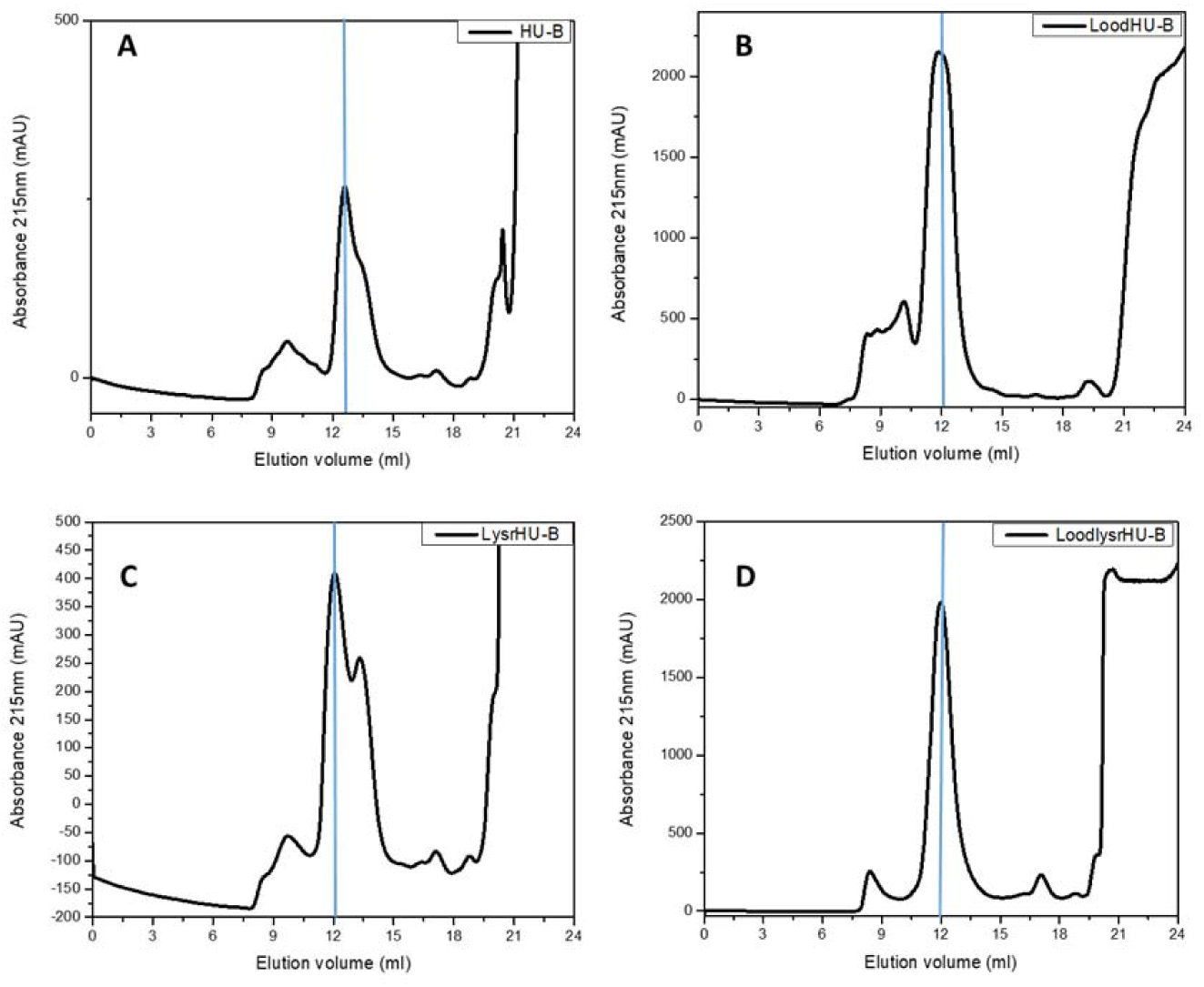
Gel filtration chromatograms of HU and its DNA-binding site-ablated variants. **a**, HU-B. **b**, LoodHU-B. **c**, LysrHU-B. **d**, LoodLysrHU-B.

## Notes

### Competing Interest Statement

The authors have declared no competing interest.

